# Spatial differences in elemental stoichiometry and essential fatty acid content of food sources and consumers in a stream food web

**DOI:** 10.1101/2023.06.14.544725

**Authors:** Monica Torres-Ruiz, John D. Wehr, Alissa A. Perrone

**Affiliations:** The Louis Calder Center – Biological Field Station and Department of Biological Sciences, Fordham University, Armonk, NY 10504 USA; National Center for Environmental Health (CNSA), Institute of Health Carlos III, Majadahonda, Spain 28220

**Keywords:** trophic transfer, biomarker, algae, bryophyte, aquatic insect, leaf detritus, nutrient, location

## Abstract

Our study characterizes spatial differences in food webs of two reaches of a New York 3^rd^-order stream differing in light availability. Food web components were analyzed using marker fatty acids (FAs). Food source nutritional quality for consumers and predators (insect larvae) was measured through stoichiometry of C, N and P and essential FAs. There were strong imbalances between detrital food sources (low N, P and essential FAs) and insects in both shaded and open reaches, and food sources and invertebrate consumers were differentially affected by light with respect to their elemental stoichiometry and essential FA content. Biochemical patterns indicated that invertebrates fed selectively on higher-quality sources (algae, bryophytes, epiphytic bacteria, transported matter) and less on lower-quality periphyton and benthic matter. In addition to confirming the importance of autochthonous food sources for stream invertebrates, this study has highlighted that local ecological processes driven by changes in light availability and canopy-derived nutrient-rich matter can alter the micro- and macronutrient content of primary producers and detrital matter. These changes tracked differently within each consumer and varied with types of nutrient. Invertebrates exhibited a greater degree of homeostasis with respect to N and P than their essential FAs, and across trophic levels.

## Introduction

Until recently, the importance of local spatial scale has been overlooked in studies of stream and other food webs, even though it is widely recognized that spatial heterogeneity can greatly affect our understanding of how food webs affect key ecosystem processes (Berg and Bengtsson 2007; Dodds et al. 2018). Spatial variation can profoundly affect food sources through changes in energy content, community composition, and system productivity (Findlay et al. 1996; Manfrin et al. 2016). The nutritional quality of food sources for macroinvertebrate consumers is also likely to vary spatially because of differences in physical and chemical conditions within the river channel (Winemiller et al. 2010). In temperate forested streams, benthic algae are limited by light availability in shaded locations, whereas sections open to sunlight are areas of increased algal production (Hill et al. 1995). These spatial dynamics can cause changes in the nutritional quality of food sources (Cashman et al. 2013; Huggins et al. 2004; Torres-Ruiz et al. 2007), which may in turn cause dietary shifts in consumers that could alter relative importance of allochthonous vs. autochthonous food sources for stream consumers (Bing et al. 2015; Closs and Lake 1994; Finlay et al. 2002).

Changes in food quantity and quality have been shown to affect macroinvertebrate consumer production and animals tend to selectively feed on food sources that provide them with essential nutrients and/or increase their omnivory rates (Demi et al. 2018; Guo et al. 2018; Lancaster et al. 2005; Sabater et al. 2011; Zhang et al. 2018). Food source nutrient content and stoichiometry within a stream could vary depending on location, but this has seldom been studied in any detail (but see Tsoi et al. 2011). With regard to primary producers, light availability has been shown to alter the elemental stoichiometry of epilithic algae, increasing C:nutrient ratios, due to increased rates of C fixation relative to nutrient availability (Dickman et al. 2006; Sterner and Elser 2002; Sterner et al. 1997). However, the opposite has also been shown under conditions of photoinhibition when exposed to high light conditions, causing lower C∶nutrient ratios in periphyton and herbivorous consumers (Martyniuk et al. 2019). Consumer stoichiometry is not always homeostatic. It is possible that C:nutrient ratios in food may be affected by differences in location, and thereby influence invertebrate C:nutrient ratios. Such an effect could result from dietary shifts and/or adjustment of body stoichiometry to that of specific food items, especially among trophic specialists (Halvorson et al. 2019; Liess and Hillebrand 2006; Persson et al. 2010).

In addition, location could also affect food source content of other essential nutrients such as polyunsaturated fatty acids (PUFAs) 20:4ω6, 20:5ω3, which are prominent in aquatic invertebrates and have been shown to be critical for their growth and reproduction (Cashman et al. 2016; Guo et al. 2018; Guo et al. 2016a; Sushchik et al. 2003; Torres-Ruiz et al. 2007; Torres-Ruiz et al. 2010; Twining et al. 2016). These essential FAs are more prominent in aquatic primary producers such as algae and bryophytes but are very scarce or even absent in sources derived from terrestrial inputs (Cashman et al. 2016; Guo et al. 2018; Guo et al. 2016b; Guo et al. 2016c; Torres-Ruiz and Wehr 2020; Torres-Ruiz et al. 2007; Torres-Ruiz et al. 2010). Spatial differences in light availability and temperature have been shown to affect periphyton PUFA content. For example, increased light availability resulted in lower levels of essential 20:5ω3 and 22:6ω3 in stream periphyton, but greater levels of 18C fatty acids (Cashman et al. 2013; Guo et al. 2016a). For a single algal or bryophyte species in a stream, spatial differences in light or temperature should affect PUFA content, but few studies have tested this idea (but see Liu et al. 2016). FA content of allochthonous food sources could also be affected by different light regimes due to changes in microbial-algal community structure (Franken et al. 2005; Guo et al. 2016d). All these spatial changes in primary producer C:nutrient ratios and FA content could in turn affect consumer food choice, consumer FA profiles, and ultimately how energy is transferred along the food chain.

In recent years, chemical tracers such as fatty acids and stable isotopes have been used to study stream food webs (Cashman et al. 2016; Guo et al. 2017; Torres-Ruiz and Wehr 2020; Torres-Ruiz et al. 2007; Twining et al. 2017). For example, fatty acids such as 16:1ω7 and 20:5ω3, 20:4ω6 as well as different 16 Carbon PUFAs, and a high (> 1) ratio of all omega-3 to omega-6 FA ([Σω3 FAs)/(Σω6 FAs]) are markers algal and bryophyte matter (Desvilettes et al. 1994; Hill et al. 2011; Napolitano et al. 1997; Torres-Ruiz and Wehr 2020; Torres-Ruiz et al. 2007; Torres-Ruiz et al. 2010). On the other hand, bacterial matter is rich in 16:1ω7, 18:1ω7 and odd-chain FAs (Kharlamenko et al. 1995; Taube et al. 2018; Torres-Ruiz and Wehr 2010), and detrital matter is rich in long chain saturated FAs (Torres-Ruiz and Wehr 2010). However, few studies combine the use of multiple chemical tracers of assimilated food items, such as fatty acids and other factors affecting energy transfer in streams, such as elemental stoichiometry (but see Twining et al. 2017; Volk and Kiffney 2012). Our previous work (Torres-Ruiz and Wehr 2020) has shown how stoichiometry and fatty acid data can complement each other in order to better understand a stream food web. In the present study, we examine these complementary data in contrasting spatial sections of a small river. Our primary goal was to analyze stoichiometry and fatty acid content of primary producers and consumers (herbivores and predators) in two locations within a stream with contrasting light regimes. We aim to investigate differences in consumer FA and C:nutrient content and how these are potentially affected by food selection, driven in turn by differences in the nutritional quality of available food sources. We hypothesize autochthonous and allochthonous matter PUFAs and elemental nutrients (C, N, P) will vary with light availably and these changes will be reflected in consumers. We will examine (1) the nutritional quality of food sources based on C, N, and P stoichiometry and essential fatty acids, (2) the possible relationships between these variables, and (3) trophic links using fatty acid markers of assimilated food.

## Materials and Methods

### Field Procedures

Two 40-m reaches of a 3^rd^-order stream, the Cross River (Westchester, County NY, USA), were sampled: (1) a shaded reach (CRS) with 95% canopy cover (mixed deciduous and conifer) located at N 41°15.732, W 73°35.808, and (3) an open canopy reach (CRO) with 5% canopy cover which runs through grassland (N 41°15.546, W 73°34.632). The two reaches are located in a protected nature preserve within 2 km of each other. Flow conditions in both sites were similar: riffle (60%) and run (30%), with a few pools (10%). Physicochemical conditions on the sampling date were otherwise similar in both reaches (means shown): width = 6.7 m; depth = 25 cm; pH = 7.6; NH_4_^+^ + NO_3_^-^ = 146 μg N/L; PO_4_^3-^ = 10.5 μg/L; Temperature = 18.5°C.

Collected food sources were operationally defined and sampled as described previously (Torres-Ruiz et al., 2019). Briefly, allochthonous food sources (terrestrial matter) were defined as benthic organic matter (BOM), and transported particulate organic matter (TOM). Five replicates of each were collected by suspension of a 20 μm mesh plankton net for 5 minutes (TOM) or from pools using a petri dish to core a depth of 2 cm (BOM). In addition, 5 replicates of leaf detritus from pools were collected from each reach in zip-lock bags, leaves collected from each site were visually different, with mostly freshly fallen green leaves collected at the shaded section of the river, compared to mostly decomposed leaves at the open section.

Five replicates of mixed periphyton, filamentous algae, and bryophytes were collected from each reach. Film covered cobles of 15–30 cm were collected for mixed periphyton whereas filamentous algae, and bryophytes (single species) were collected separately by hand from five boulders and washed to remove sediment and invertebrates. A 243-μm-mesh Surber sampler was used to collect macroinvertebrates from 5 random locations within each reach. Invertebrates were rinsed into a tray and placed in separate 50 mL tubes with stream water. Animals of each taxon were carefully chosen to be of similar developmental stage (similar size and wing pad length). All samples were placed in zip-lock plastic bags, stored on ice, and returned to the laboratory for processing within 2 h of collection.

### Laboratory Processing

Laboratory processing was done as described previously (Torres-Ruiz and Wehr 2020). Briefly, we divided TOM and BOM into three size fractions using two nested sieves (1 mm and 250 μm mesh) to separate coarse (C: > 1 mm) and fine (F: 250 μm to ≤ 1mm) fractions; ultrafine (UF: < 250 μm) was defined as matter that passed through the 250 μm sieve but was retained on a Whatman GF/F glass fiber filter (GE/Whatman, U.K.). CTOM, CBOM, and leaves were gently rinsed with deionized water (dH_2_O) and blended (≤ 5s) before filtration. Periphyton covered cobles were scraped using brushes and dH_2_O. Pre-ashed Whatman GF/F filters were used for dry mass (DM), nutrient content (Carbon, Nitrogen, Phosphorus), and FA analyses. Filters for FA extraction were stored under N_2_ and frozen (–20°C), and those for nutrients and dry mass quantification were dried in an oven (80°C) for later analysis.

Bryophytes and filamentous algae were identified to genus and species. The main autochthonous primary producers occurring in both reaches of the Cross River were the bryophyte *Fontinalis dalecarlica* Bruch & Schimp. (hereafter *Fontinalis*); the red alga *Lemanea fluviatilis* (Linnaeus) C.Agardh, 1811 (*Lemanea*); the green alga *Cladophora glomerata* (Linnaeus) Kützing, 1843 (*Cladophora*), and mixed periphyton. In addition, a third macroalgal species, the xanthophyte *Vaucheria* sp. A.P. de Candolle (*Vaucheria*), occurred only in CRO. These algae/bryophytes were rinsed thoroughly with dH_2_O until visibly clean and tips (top 2 cm) were cut and separated into three different portions (Wehr et al. 1983). Material for FA analysis was weighed, placed in a 15 mL tube under N_2_ and frozen (–20° C); material for DM and nutrient analysis was weighed, placed in two tin cups and dried (80° C). Macroinvertebrates were quickly sorted to genus or family and allowed 20- to 30-h gut-clearance in clean stream water (at 4° C). Following this time macroinvertebrates were clearly separated by species using a dissecting microscope.

Larvae of nine macroinvertebrate taxa (all aquatic insects) were sufficiently abundant in both sections of the river to complete both sets of chemical analyses required for the present study: *Hydropsyche* sp. (collector, Trichoptera); *Epeorus* sp., and *Stenonema* sp. (grazers, Ephemeroptera); *Isonychia* sp., (collector, Ephemeroptera); *Glossosoma* sp. (scraper, Trichoptera); *Psephenus* sp. (scraper, Coleoptera); *Rhyacophila* sp. (predator, Trichoptera), *Paragnetina* sp. (predator, Plecoptera), and *Nigronia* sp. (predator, Megaloptera). Three groups of each species (several animals of similar size/developmental stage) were gathered and weighed. One group was placed under N_2_ and frozen (–20° C) for FA extraction and the rest were placed in tin cups and dried (80° C) for DM and elemental nutrient analyses.

Samples to be analyzed for total Carbon (C), and total Nitrogen (N) were homogenized to a fine powder (< 1 mm), placed in a pre-weighed tin capsule, weighed using a Perkin Elmer AD-6 ultra-microbalance (Perkin-Elmer, Waltham, MA) (nearest 0.01 mg) and sent to the University of California Davis for analysis. C and N were analyzed by dry combustion (Europa Hydra 20/20 analyzer, Northwich, UK). A modified method of Solorzano & Sharp (1980) was used to analyze particulate Phosphorus (P). Total P in dried samples was extracted, converted to orthophosphate (PO_4_^3-^), analyzed using antimony-molybdate, and measured spectrophotometrically (Shimadzu UV-160, Kyoto, Japan). Fatty acids in food sources and consumers were analyzed as completed in prior studies (Torres-Ruiz and Wehr 2020). Briefly, lipids were extracted in chloroform/methanol (2:1v/v), methylated with BF_3_ (10–15% w/v in methanol), resuspended in hexane, and concentrated under N_2_. A Hewlett-Packard 5890 gas chromatograph (Hewlett-Packard, Avondale, PA) fitted with a Supelco Omegawax 320 capillary column (30 m x 0.32 mm; Supelco®, Bellefonte, PA, USA) and an FID detector was used to separate and analyze FA methyl esters. Individual FAs were identified using certified standard mixtures (SupelcoTM 37 Component FAME mix, Menhaden Oil: PUFA-3, and Supelco bacterial acid methyl esters mix [BAME]), and a few single FAME standards. Reagent blanks were processed and analyzed alongside samples.

### Data Analyses

Nonmetric multidimensional scaling (NMDS) was used to study similarities of FA profiles between potential food sources and macroinvertebrates. NMDS ordinations were done on Bray– Curtis distances using PC-ORD (multivariate analysis of ecological data, version 4.0; MjM Software, Gleneden Beach, Oregon). FA variables used in these analyses were: (1) essential fatty acids (EFAs: 18:2ω6, 20:4ω6, 18:3ω3, 20:5ω3); (2) sums of saturated FAs (SAFA), monounsaturated FAs, and PUFAs; (3) potential biomarker FAs, including diatom markers 16:1ω7 and 16:2ω4, bacterial markers 18:1ω7 and the sum of odd-chain FAs and fatty acids identified through the use of BAME standard (odd + BAME), and (4) the ratio Σω3/Σω6. Proportion of variation explained by each axis was calculated and correlations were performed between NMDS axis loadings and FAs (Torres-Ruiz and Wehr 2020).

Percent PUFAs (18:2ω6, 18:3ω3, 20:5ω3 and 20:4ω6), Σω3/Σω6, C/N, C/P, and N/P ratios, were compared for each food source and consumer category between sites using two-tailed Student *t*-tests. Data were tested for normality and homogeneity of variance assumptions and transformed where required. Percent data were arcsine transformed and concentration data were square-root transformed (Sokal and Rohlf 1995). We used a ratio Σω3/Σω6 significantly greater than 1 (tested by a one-sample Student’s *t*-test) to indicate relative importance of autochthonous vs. allochthonous origin of a food source (Desvilettes et al. 1994; Torres-Ruiz et al. 2007). All reported FA values represent untransformed means. An *a priori* probability for a type I error of ≤ 0.05 was used throughout. Statistical analyses were performed using Systat v. 11.

Our interpretation of nutrient data has assumed invertebrate C, P, and N homeostasis, and did not take into consideration consumer growth rates (Frost et al. 2006; Small and Pringle 2010). Likewise, we have assumed whole body fatty acid/nutrient content reflects diet signatures as has been shown before (Guo et al. 2016a; Harlıoğlu et al. 2015; Torres-Ruiz et al. 2010).

## Results

### Fatty acids

General similarities in FA composition of between consumers and food sources were summarized using non-metric multidimensional scaling (NMDS; Fig. 1). NMDS axes 1, 2 and 3 explained 28.8%, 27.4% and 25.4%, of the total variation in the dataset of the open reach (CRO: Fig. 1A, B) and 31.1%, 28.7% and 20.5% respectively for the shaded reach (CRS: Fig. 1C, D). Both analyses revealed consistent patterns of FA signatures were among replicates for each of the trophic levels and food categories.

**Fig. 1.**
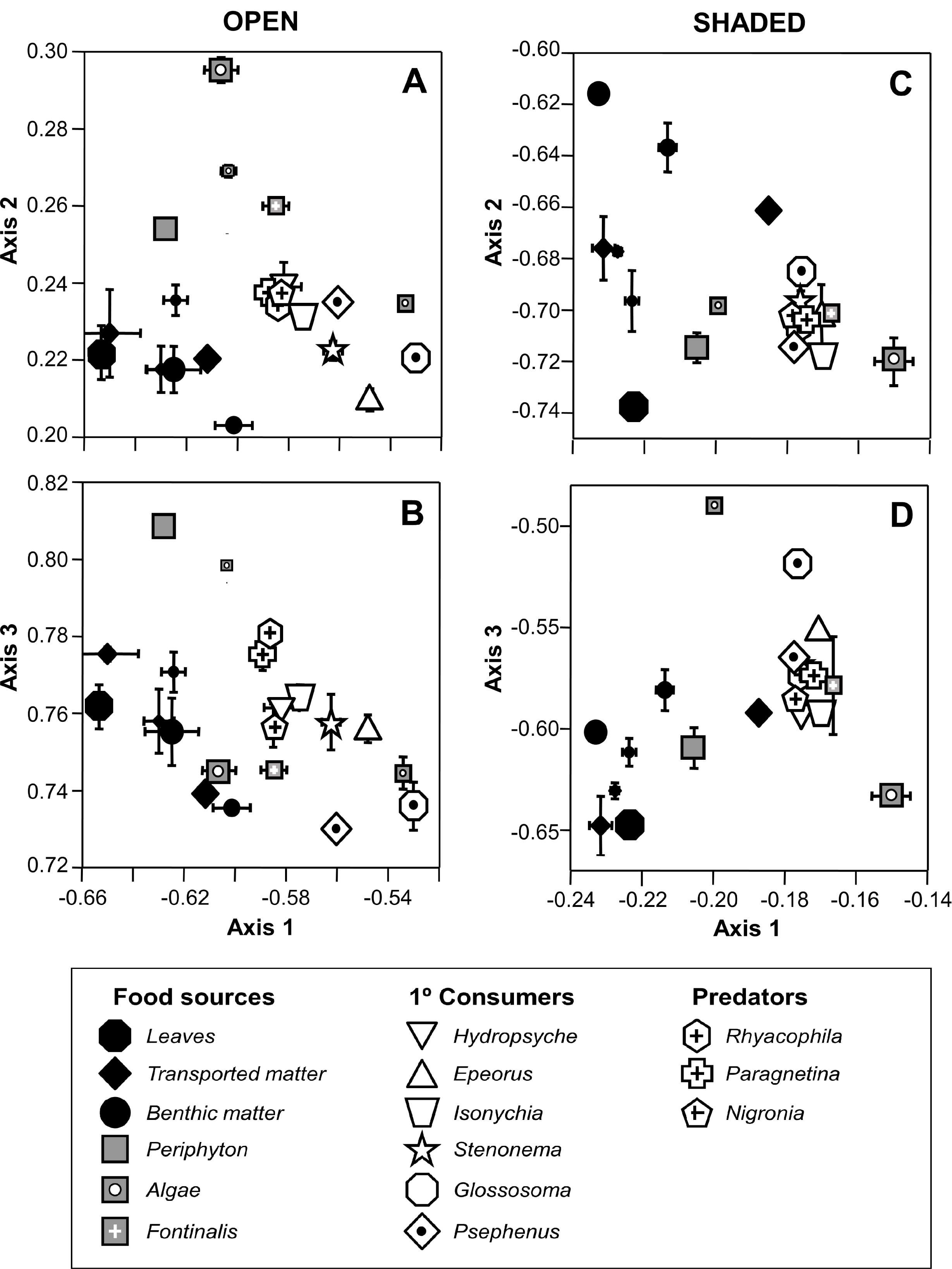
Nonmetric multidimensional scaling plots of food categories (black = allochthonous, grey = autochthonous) and insect larvae (open symbols) in the open (**A** and **B**) and shaded (**C** and **D**) sections of the Cross River, based on their fatty acid content (values represent mean loadings (±1 SE). Large black diamond = coarse transported matter; medium black diamond = fine transported matter; small black diamond = ultrafine transported matter; large black circle = coarse benthic matter; medium black circle = fine benthic matter; small black circle = ultrafine benthic matter; large white-dotted grey square = *Lemanea*; medium white-dotted grey square = *Cladophora*; small white-dotted grey square = *Vaucheria*; small white-crossed grey square = *Fontinalis*.

In the open reach (CRO), consumer positions were widely dispersed, with *Epeorus, Stenonema, Psephenus,* and especially *Glossosoma* plotting closest to the green alga *Cladophora* (Fig. 1A, B). These animals were characterized by PUFAs that correlated positively with axis 1 (PUFA (*r* = 0.8), 18:3ω3 (*r* = 0.7), 16C-PUFAa&b (*r* = 0.6), 20:5ω3 (*r* = 0.6), and Σω3/Σω6 (*r* = 0.6), [*P* ≤ 0.001 for all]). *Rhyacophila* and *Paragnetina* plotted centrally, while *Hydropsyche, Isonychia* and *Nigronia* separated along axis 3, closer to the moss *Fontinalis* (Fig. 1B). Most types of allochthonous matter (leaves, benthic matter, and transported matter) plotted low on axes 1, 2 and 3 (Fig. 1A, B). This matter was characterized by FAs correlated negatively with axis 1 [SAFA (*r* = −0.8), 22:0 (*r* = −0.6), 20:0 (*r* = −0.4), and 18:2ω6 (*r* = −0.3)]; axis 2 [(17:0 (*r* = −0.6), 20:0 (*r* = −0.5), 22:0 (*r* = −0.5), and 18:1ω7 (*r* = −0.5)] and axis 3 [(20:2 (*r* = −0.5), 17:1 (*r* = −0.4), bacterial FAs (*r* = −0.4), and 16:1ω9 (*r* = −0.4)]; *P* ≤ 0.001 for all.

In the shaded reach (CRS), most insect larvae plotted in a tight group close to (in order of closeness): the aquatic moss *Fontinalis*, CTOM, the green alga *Cladophora,* mixed periphyton, and the red alga *Lemanea* (Fig. 1C, D). This insect group was characterized by typical algal polyunsaturated FAs (PUFAs), which correlated positively with axis 1 [(20:5ω3 (*r* = 0.8), 20:4ω6 (*r* = 0.7), 16C-PUFAa (*r* = 0.5) and total PUFA (*r* = 0.6)]; *P* ≤ 0.001). It was also characterized by FAs that correlated negatively with axis 2 [20:5ω3(r = −0.4), 18:1ω9 (*r* = −0.3), 18:3ω3 (*r* = −0.3), and total PUFA (*r* = −0.4)]; *P* ≤ 0.01), and by algal marker FAs that correlated positively with axis 3 [16C-PUFAb (*r* = 0.8), 16:3ω4 (*r* = 0.7), 16C-PUFAa (*r* = 0.6), 18:3ω3 (*r* = 0.5), and total PUFA (*r* = 0.6)]; *P* ≤ 0.001. The scraper-grazer *Glossosoma* was plotted distant from other consumers along axis 3, but closer to *Cladophora.* Most types of allochthonous matter plotted in a region low on axes 1 and 3 (Fig. 1D) characterized by saturated FAs (SAFAs) and bacterial markers, and which correlated negatively with axis 1 [22:0 (*r* = −0.7), 24:0 (*r* = - 0.6), 20:0 (*r* = −0.6), BAME (*r* = −0.6)]; *P* ≤ 0.001. However, they were more dispersed along axis 2, with CBOM and FBOM plotting in a region correlated with long chain saturated FAs and bacterial FAs [(24:0 (*r* = 0.8), 20:0 (*r* = 0.7), 22:0 (*r* = 0.6), and bacterial markers (r = 0.6)], and green leaves and UFBOM plotting low on axis 2 which correlated with higher omega-3 PUFA content (see above).

Comparing essential FA profiles in open and shaded sites revealed spatial differences in most food-source categories and animals (Figs. 2-4). EFAs 18:ω6 and 18:ω3 were less abundant in smaller size categories of BOM and POM at both sites (Fig. 2A and B). In addition, bacterial FAs were more abundant in smaller size categories of BOM but no differences were found with site (Fig. S1A). Leaves, UFBOM and POM had higher 18:ω6 and 18:ω3 content at CRS, possibly due to green leaves, pollen, and other material fallen from the summer canopy. Longer and typical algal EFA 20:5ω3, as well as the Σω3/Σω6 ratio, were less in BOM at the shaded section (Figs. 2A and 4A), although this difference was only significant for both in the UFBOM fraction. Content of EFA 20:4ω6 was very small in both BOM and POM at both sites (Fig. 2A and B).

**Fig. 2.**
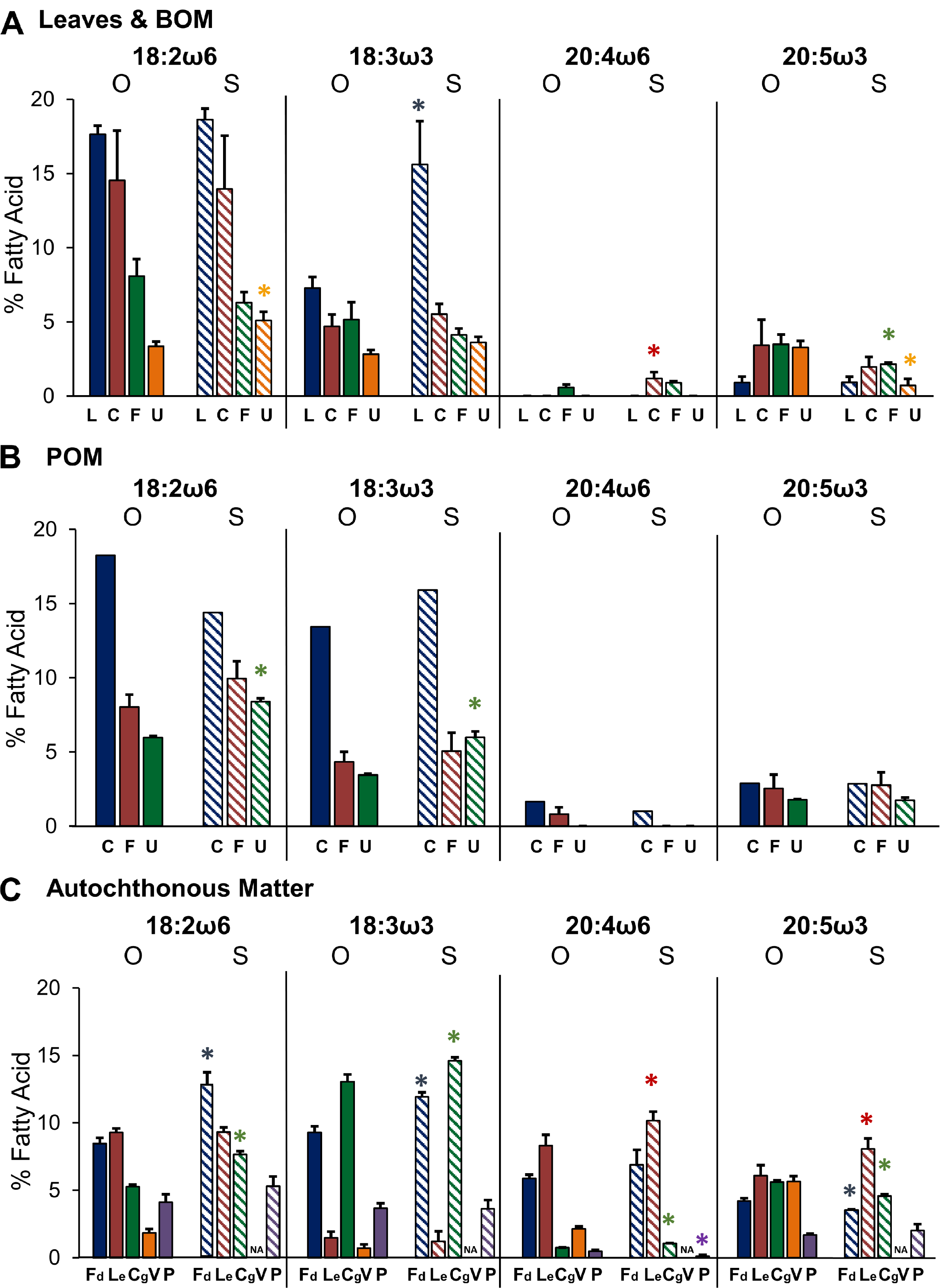
Changes with site of essential fatty acid content of leaves and benthic matter (**A**), particulate matter (**B**) and autochthonous matter (**C**) in the open (O; solid bars) and shaded (S; striped bars) areas of the Cross River. Values represent means (±1 SE). Asterisk above bars represent significant differences with site (p < 0.05). L = leaves; C = coarse; F = fine; U = ultrafine; Fd = *Fontinalis*; Le = *Lemanea*; Cg = *Cladophora*; V = *Vaucheria*; P = periphyton.

**Fig. 3.**
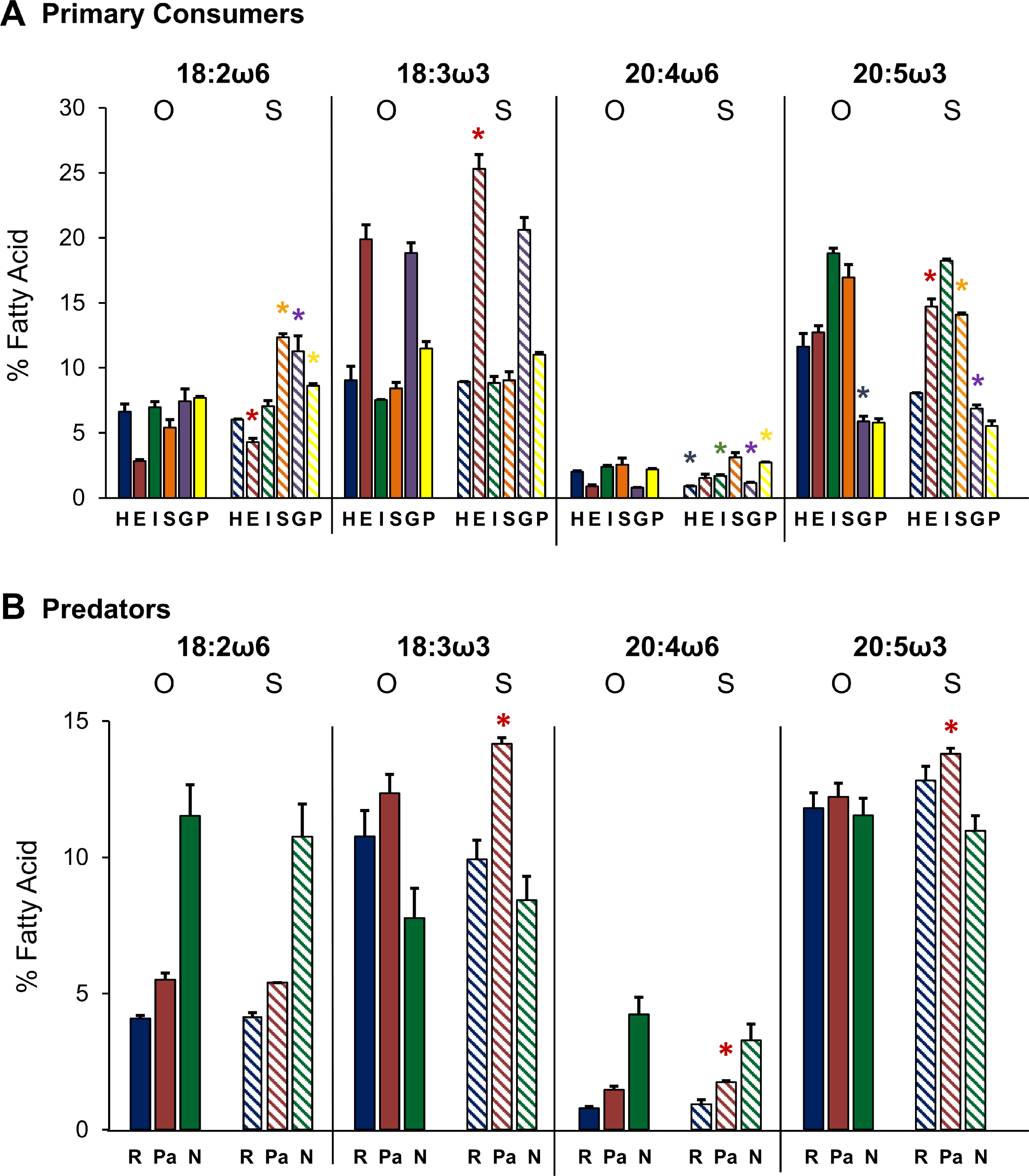
Changes with site of essential fatty acid content of primary consumers (**A**) and predators (**B**) in the open (O; solid bars) and shaded (S; striped bars) areas of the Cross River. Values represent means (±1 SE). Asterisk above bars represent significant differences with site (p < 0.05). H = *Hydropsyche*; E = *Epeorus*; I = *Isonychia*; S = *Stenonema*; G = *Glossosoma*; P = *Psephenus*; R = *Rhyacophila*; Pa = *Paragnetina*; N = *Nigronia*.

**Fig. 4.**
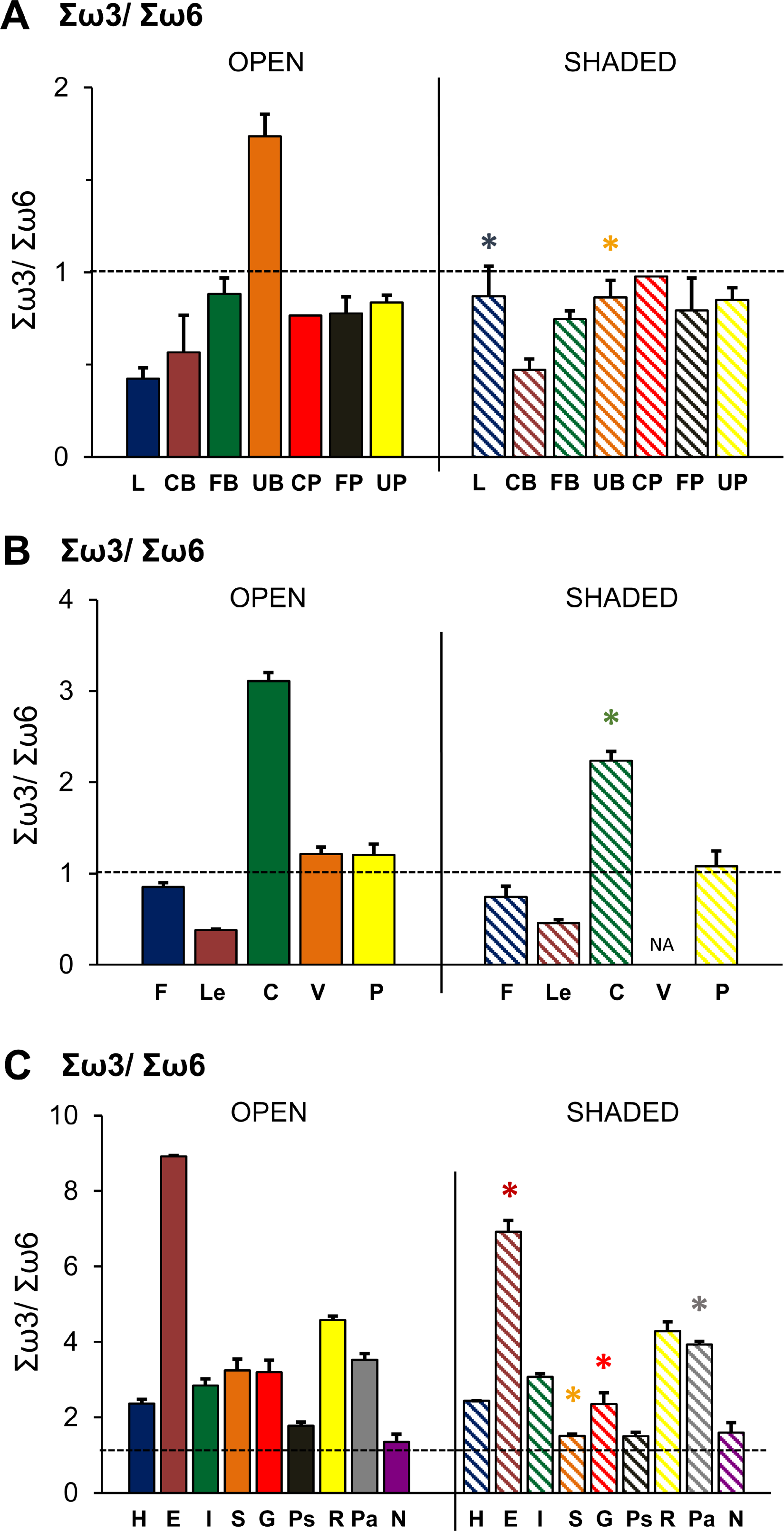
Changes with site in the Σω3/Σω6 ratio of benthic and particulate matter (**A**), autochthonous sources (**B**), and animals (**C**) in the open (O; solid bars) and shaded (S; striped bars) areas of the Cross River. Values represent means (±1 SE). Asterisk above bars represent significant differences with site (p < 0.05). L = leaves; CB = coarse benthic; FB = fine benthic; UB = ultrafine benthic; CP = coarse particulate; FP = fine particulate; UP = ultrafine particulate; F = *Fontinalis*; Le = *Lemanea*; C = *Cladophora*; V = *Vaucheria*; P = periphyton; H = *Hydropsyche*; E = *Epeorus*; I = *Isonychia*; S = *Stenonema*; G = *Glossosoma*; P = *Psephenus*; R = *Rhyacophila*; Pa = *Paragnetina*; N = *Nigronia*.

All autochthonous food source categories differed in their EFA content between sites, although trends depended on the species of autotroph (Fig. 2C). In general, EFAs 18:ω6 and 18:ω3 were abundant in the moss *Fontinalis* and the alga *Cladophora,* and were significantly greater at the shaded site. The richest sources of 20:4ω6 at both sites were *Fontinalis* and the red algae *Lemanea*. This particular FA was elevated in most autochthonous sources at the shaded site, although only significantly in *Lemanea* and *Cladophora*. The red algae *Lemanea* was the richest 20:5ω3 source and it increased significantly at the shaded site. Contrarily, *Fontinalis* and *Cladophora* decreased their 20:5ω3 content at the shaded site. Mixed periphyton had similar EFA content at both sites, and this relatively stable property was also true for saturated, monounsaturated, and bacterial FAs (Fig. S1C). Periphyton had high content of MUFAs, SAFAs and BAFAs (Fig. S1C), and low 20:5ω3 (Fig. 2C).

Differences in EFA content of primary consumers between sites depended on the invertebrate species and category of FA (Fig. 3A). The grazing caddisfly *Glossosoma* varied most between sites in all EFAs (although non-significant for 18:3ω3). Net-spinning *Hydropsyche* had significantly less 20:4ω6 and 20:5ω3 at the shaded site (Fig. 3A), together with greater content of saturated FAs (Fig. S2A). *Epeorus* and *Glossosoma* had, in general, greater EFA content at the shaded site, although not all differences were statistically significant (Fig. 3A). In addition, these two animals also had lower MUFA and SAFA content at the shaded site (Fig. S2A).

EFA content in the collector *Isonychia* was the least variable, with changes only in 20:4ω6 (less at shaded site), while the grazer *Stenonema* had significantly greater 18:2ω6 and significantly lower 20:5ω3 content at the shaded site. Another grazer category, *Psephenus* had greater content of both ω6 EFAs at the shaded site. The only predator that changed EFA profile with site was *Paragnetina*, with higher EFA content at CRS (Fig. 3B). This animal also had significantly lower SAFA content at this site (Fig. S2B). All other predators maintained similar EFA profiles at the two sites (Fig. 3B).

The Σω3/Σω6 ratio was inversely related to particle size in BOM, being greatest in UFBOM at both sites (Fig. 4A). UFBOM had an Σω3/Σω6 ratio > 1.0 at the open site but was < 1.0 at the shaded site. Aside from UFBOM and green leaves (shaded site), all other benthic and transported matter had similar Σω3/Σω6 ratios at both sites, all significantly less than 1.0 (*P* < 0.05). *Fontinalis dalecarlica* and *Lemanea* also had Σω3/Σω6 ratios lower than one and were similar at both sites, in spite of changes in individual EFAs (Fig. 4B); ratios were due to their greater 18:2ω6 and 20:4ω6 content versus ω3 FAs. In contrast, the green algae *Cladophora* (Fig. 2C) had the highest Σω3/Σω6 ratio at both sites, although it was significantly less at the shaded section. However, these three food sources had similar 20:5ω3 content and were, at this time of year, very rich EFA sources for invertebrates. Periphyton Σω3/Σω6 ratios were similar at the two locations and not significantly different from 1.0 (Fig. 4B). All animals, except for *Nigronia* (both sites) and *Psephenus* (shaded site) had Σω3/Σω6 ratios significantly greater than 1.0 at both sites (*P* < 0.05) (Fig. 4C). A significant decrease in this ratio at the shaded site was observed in the mayflies *Epeorus* and *Stenonema* and the caddisfly *Glossosoma*. In the case of *Epeorus* and *Glossosoma* at the shaded site, ratios were affected by a greater increase in ω6 FAs than increases in ω3 FAs (Fig. 3A). In *Stenonema* at the shaded site, the shift was attributed to an increase in 18:2ω6 and a decrease in 20:5ω3. Although non-significant, the Σω3/Σω6 of *Psephenus* was also less at the shaded site (Fig. 4C), which was due to an increase in 18:2ω6 and 20:4ω6 at the shaded site together with similar ω3FA content at both sites. The predator *Paragnetina* had a greater Σω3/Σω6 ratio at the shaded site, in part due to larger increases in ω3 FAs greater versus increases in 20:4ω6 (Fig. 3B).

### Elemental Stoichiometry

Spatial differences in C, N and P profiles between the open and shaded sites were evident in several food source categories and consumers (Figs. 5, 6). Nutrient content varied also with size of allochthonous matter categories, and among species of primary producers and animals. In contrast to their relatively stable FA content, elemental nutrients in periphyton and FPOM were quite variable. Specifically, small size categories of allochthonous matter had much lower C/N and C/P ratios than larger fractions at both sites (e.g. CBOM C/P = 2782 vs UFBOM C/P = 271 at CRO; Fig. 5 A, C). This was associated with greater C content in leaves and coarse particulate matter fractions (Fig. 6). Allochthonous sources had in general lower N and P content than autochthonous sources (Fig. 6), although their C/N and C/P ratios were sometimes similar (e.g. *Cladophora* C/P = 280 vs. UFPOM C/P = 275 at CRS; Fig. 5 A,C). This was especially true of ultrafine fractions, mainly due to their lower C content (e.g. UFPOM C = 175 mg/g vs. *Cladophora* C = 282 mg/g), and that autochthonous matter had generally higher P content (Fig. 6). Although N content of autumn-shed leaves was overall low, concentrations were significantly (40%) greater at the shaded site, while P content was largely unchanged (Fig. 6). Elemental stoichiometry of transported matter (FPOM) differed greatly (10 to 100-fold) with site (e.g. 399 mg/g C and 3.8 mg/g P in CRS versus 186 mg/g C and 0.8 mg/g P in CRO, Fig. 6B. D). This was reflected in a significant decrease in C/P and N/P in CRS (FPOM C/P = 170 in CRS vs. 250 in CRO, *P* = 0.03; Fig. 5B, E). On the other hand, its C/N ratio was greater at CRS and associated with greater increase in Carbon than in N (Fig. 6). A similar trend was observed for UFPOM (Fig. 5C).

**Fig. 5.**
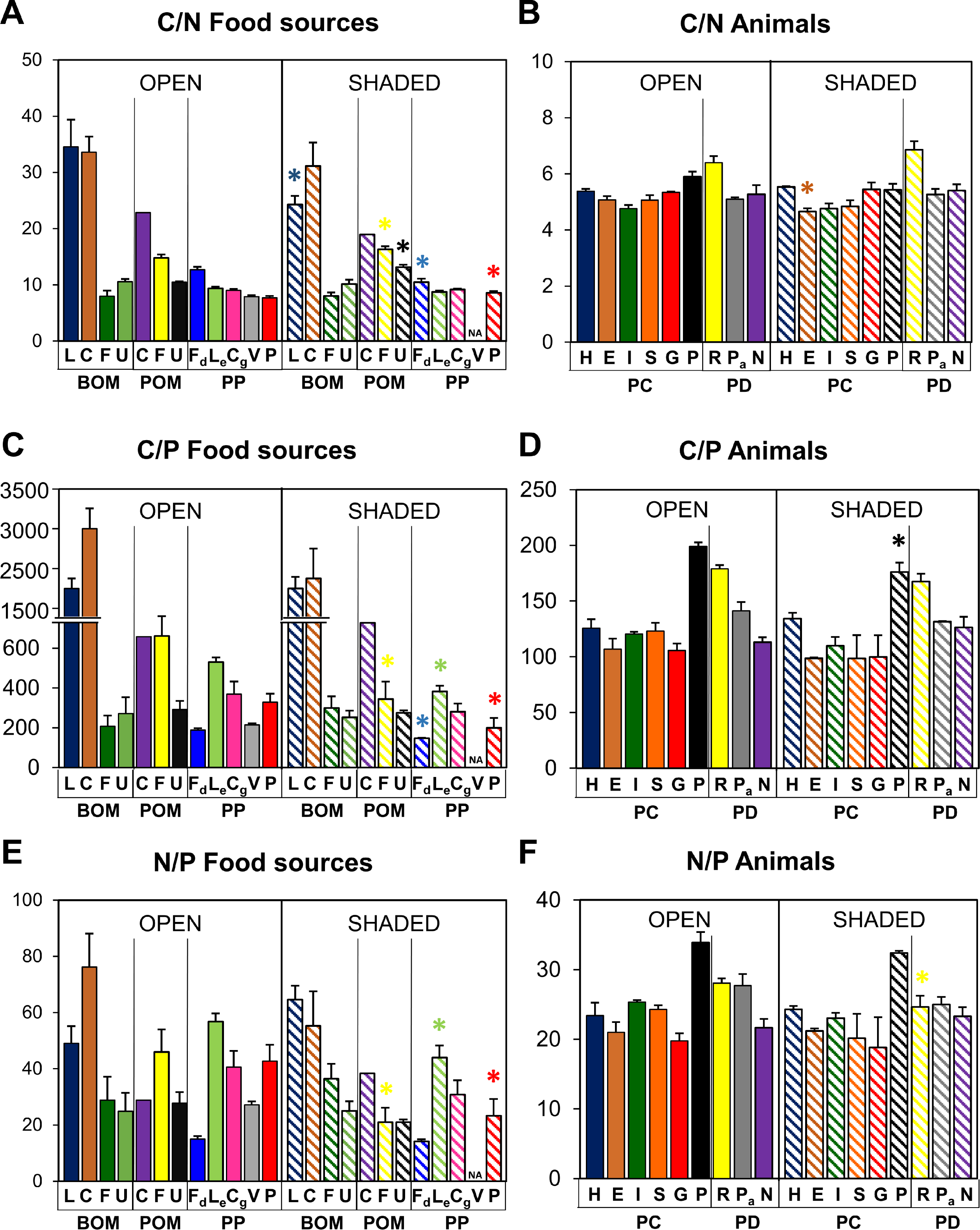
Changes with site in the C/N, C/P and N/P ratios of food sources (**A, C, E** respectively), and animals (**B, D, F**) in the open (O; solid bars) and shaded (S; striped bars) areas of the Cross River. Values represent means (±1 SE). Asterisk above bars represent significant differences with site (p < 0.05). Note axis break in food source C/P. BOM = benthic organic matter; POM = particulate organic matter; PP = primary producers; PC = primary consumers; PD = predators; L = leaves; C = coarse; F = fine; U = ultrafine; Fd = *Fontinalis*; Le = *Lemanea*; Cg = *Cladophora*; V = *Vaucheria*; P = periphyton; H = *Hydropsyche*; E = *Epeorus*; I = *Isonychia*; S = *Stenonema*; G = *Glossosoma*; P = *Psephenus*; R = *Rhyacophila*; Pa = *Paragnetina*; N = *Nigronia*.

**Fig. 6.**
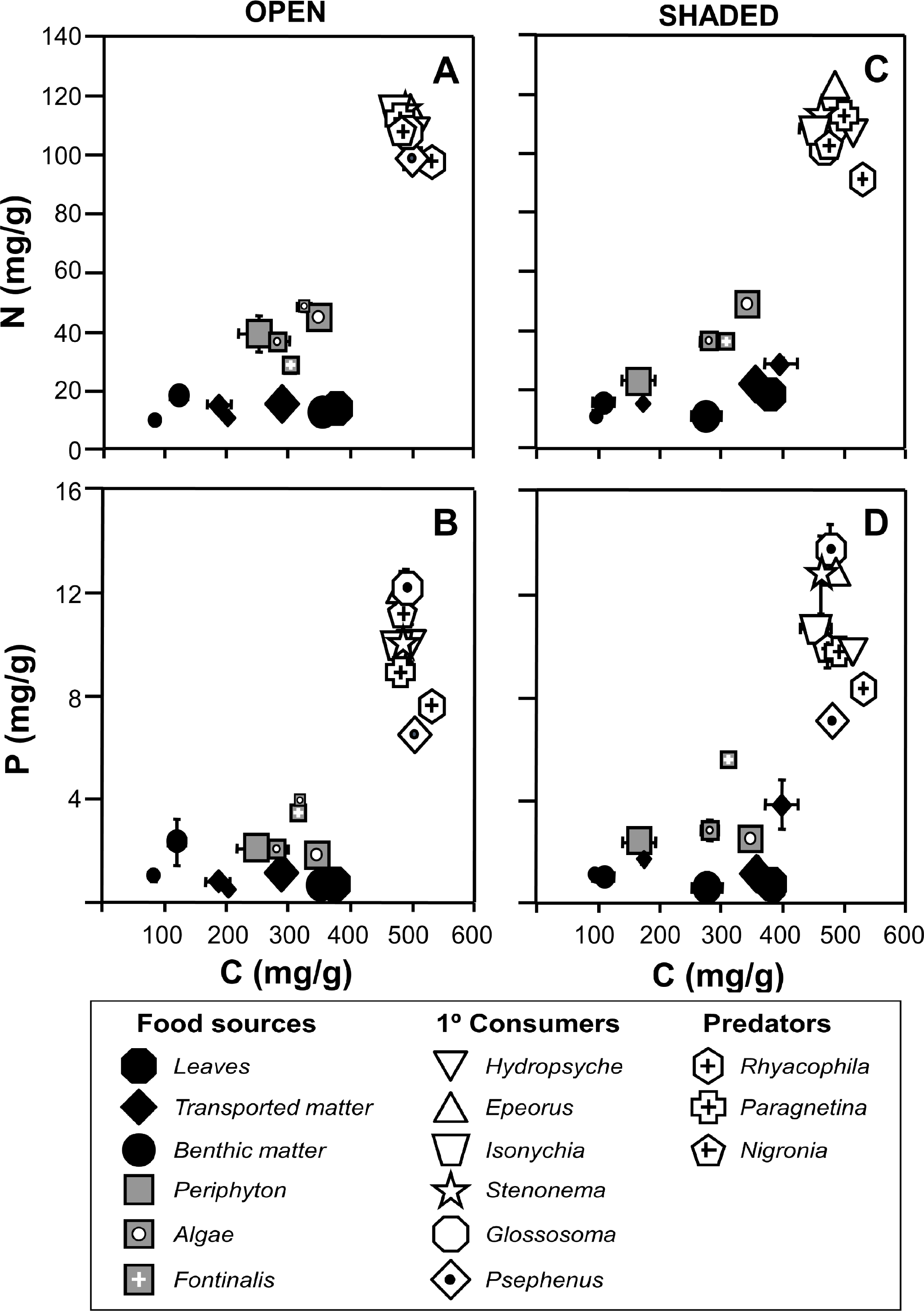
Nutrient concentrations of food categories and insect larvae of the open (**A** and **B**) and shaded (**C** and **D**) sections of the Cross River. Values represent means (±1 SE) (N = Nitrogen [mg/g dry mass], P = Phosphorus [mg/g DM], C = Carbon [mg/g DM]). Large black diamond = coarse transported matter; medium black diamond = fine transported matter; small black diamond = ultrafine transported matter; large black circle = coarse benthic matter; medium black circle = fine benthic matter; small black circle = ultrafine benthic matter; large white-dotted grey square = *Lemanea*; medium white-dotted grey square = *Cladophora*; small white-dotted grey square = *Vaucheria*; small white-crossed grey square = *Fontinalis*.

Autochthonous matter varied in nutrient content depending on species and in some cases with location. The red alga *Lemanea* had the greatest N content at CRS (46 mg/g), while the xanthophyte *Vaucheria* was the richest source in CRO (48 mg/g; Fig. 6A, C). The moss *Fontinalis* was the richest P source at both sites, being the only food source with a C/P similar to that of consumers (C/P = 188 and 146 at CRO and CRS respectively; Fig. 5C). Among autotrophs, *Fontinalis*, *Lemanea* and periphyton varied most with site. Specifically, *Fontinalis* and *Lemanea* had significantly greater N and P contents in the shaded site (e.g. *Fontinalis* P = 4.1 at CRO vs 5.5 at CRS; *P* = 0.0001; Fig. 6B, D), resulting in lower C/N and C/P ratios (Fig. 5 A, C). In contrast, mixed periphyton had lower C and N, at the shaded site, but with similar P content (Fig. 6). As a result periphyton had higher C/N ratios but lower C/P and N/P (e.g. Periphyton C/P = 70 in CRS vs. 127 in CRO, *P* = 0.04; Fig. 5A, C, E). Spatial patterns of nutrient content and stoichiometry in *Cladophora* were similar to those in *Fontinalis* and *Lemanea*, but were non-significant.

Animals varied in their nutrient content depending on species, but there were few significant differences with site. In general, consumers had lower C/N and C/P than most food sources (Fig. 5). In CRS, N content varied from 122 mg/g (*Epeorus*) to 90 mg/g (*Rhyacophila*) and in CRO from 115 mg/g (Isonychia) to 97 mg/g (*Rhyacophila*) (Fig. 6A, C). *Glossosoma* had the highest consumer P content and lowest C/P ratios in both sites (13.6 mg/g and 98 in CRS, 12.1 mg/g and 105 in CRO), while *Psephenus* the lowest P and highest C/P (7.2 mg/g and 176 in CRS, 6.5 mg/g and 198 in CRO) (Figs. 5D, 6B, 6D). Few animals varied in nutrient content and ratios with site. *Epeorus* had significantly higher N and lower C/N ratios at the shaded site (Fig. 5B), while *Psephenus* and *Rhyacophila* had higher P and lower C/P and N/P at CRS (Fig. 5D, F). *Epeorus*, *Isonychia*, *Stenonema*, *Glossosoma*, and *Paragnetina* had non-significant differences in P content between sites (Fig. 6).

## Discussion

### Spatial differences in autochthonous food quality

Our data demonstrate that the fatty acid content and elemental stoichiometry of allochthonous matter, primary producers, and consumers in local stream food webs can vary spatially between nearby reaches that differ primarily in light regime. These spatial differences have profound effects on nutrient and energy transfer within a stream.

Our data show that increased light availability at the open location had greater fatty acid and elemental stoichiometric quality of basal food sources, in part due to greater algal colonization. However, periphyton is a mixed assemblage of many species of microalgae, bacteria, fungi, and protists whose FA and elemental nutrient content can change in complex ways with differences in light (Cashman et al. 2013; Guo et al. 2016a; Hill et al. 2011; Martyniuk et al. 2019; Sanches et al. 2011). For example, Cashman et al. (2013) observed decreased 18C EFAs, but unchanged levels of 20:5ω3 in periphyton in shaded canopy conditions, whereas Guo et al. (2016a) showed an increase of 20:5ω3 and 20:4ω6. It is likely that changes in FA content of a biofilm in many studies is the result of changes in algal community composition (Steinman and McIntire 1987). In our study, periphyton was characterized by FAs typical of chlorophytes and diatoms: 16PUFAs, 18:3ω3 and 20:5ω3 (Fig. 2C, Table S1). Contrary to some previous studies, the algal FA *signature* of the periphyton assemblage in the Cross River did not change significantly with location, which suggests the algal community at the two reaches was similar, a pattern which has been seen elsewhere (Wellnitz and Rader 2003). Nevertheless, the FA signature of periphyton did vary with regard to certain specific marker FAs and key nutrient elements, which was reflected the complex mixture of organisms within this matrix. The bacterial marker FA 16:1ω7 was roughly twice that measured in the CRO river section, which received greater sunlight (12 vs. 20%; Table S1). In addition, C and N content (mg/g) of periphyton were 35% and 40% greater at CRO, which resulted in significantly lower C/N ratios (Figs. 5 and 6). FA and nutrient data taken together suggest that at the open section, there was an increase in photosynthesis, which may have also supported greater bacterial biomass. This increase in photosynthesis with light and the coupling between bacteria and algae in periphyton is consistent with the light-nutrient hypothesis, as observed elsewhere (Sanches et al. 2011; Sterner et al. 1997; Wetzel 2005). Fungi can also be a prominent part of stream periphyton (Miura and Urabe 2015). Our data show that fungal marker FAs (18:1ω9, 18:2ω6) were common in the Cross River periphyton, and occurred at greater levels in the shaded section (CRS, Fig. 2C, Table S1). In addition, N/P ratios were significantly lower at CRS (Fig. 5E). Based on our FA and N/P data, and because fungi have been shown to store P to a greater degree than N (Gulis et al. 2017), we suggest mixed periphyton in shaded habitats have a greater proportion of fungal cells. We specifically predict an order of importance within periphyton in shaded streams or streams sections as algae > fungi > bacteria, while in open streams this shifts to algae > bacteria > fungi. Our data highlight the complexity of microbial assemblages in stream periphyton (Wyatt et al. 2019), which have multiple interactions between bacteria and fungi growing on aquatic biofilms (Mille-Lindblom et al. 2006; Torres-Ruiz and Wehr 2010).

Macroalgae and the bryophyte *Fontinalis* in the Cross River had different FA and elemental nutrient content, with some differences in biochemical signatures between sites. Overall, the EFA content of these autotrophs was much greater than that of allochthonous matter and mixed periphyton, regardless of location. The green alga *Cladophora* had the highest PUFA content of all macroalgal food sources, while the xanthophyte *Vaucheria* had the least (but greater MUFA; Fig. S1C). *Cladophora glomerata* also had the highest content of ω3 PUFAs, making it the most similar, in terms of Σω3/Σω6 ratio, to consumers (Fig. 4). We know of no field studies that have documented the FA signature of *Vaucheria*, which is surprising given its commonness in many streams (Schagerl and Kerschbaumer 2009). This alga was characterized by low levels of 18:3ω3 and 18:2ω6 (1-2%), but a high percentage of 16:1ω7 (35%) and the essential 20:5ω3 at levels similar to other macroalgae (6%; Fig. 2C, Table S1). Our study also provides new data on the FA signature for the moss *Fontinalis dalecarlica*, which supplement information on another species in the genus, *F. antipyretica* (Jamieson and Reid 1976; Kalacheva et al. 2009). *Fontinalis dalecarlica* had a distinct FA signature with high levels of EFAs 18:3ω3, 18:2ω6 and 20:4ω6, but lower amounts of 20:5ω3 than macroalgae, which resulted in an Σω3/Σω6 ratio < 1.0 (Figs. 2C and 4B). Interestingly, open vs. shaded canopy conditions affected the FA signature of this moss, with lower MUFA and higher PUFA (18:3ω3, 18:2ω6; but not 20:4ω6 or 20:5ω3) when shaded (Fig. S1C). The green alga *Cladophora* exhibited similar changes with light, with higher PUFA (18:3ω3, 18:2ω6 and 20:4ω6) and lower SAFA at the shaded section (Figs. 2C and S1C). The observed changes in *Fontinalis* and *Cladophora* shown here have been observed before in *Cladophora* (Napolitano 1994), microalgae (Hu et al. 2008), other species of filamentous green algae (Liu et al. 2016).

These data suggest that greater exposure to sunlight at CRO stimulated the formation of energy-storage lipids such as MUFA (*Fontinalis*) and SAFA (*Cladophora*), whereas shaded conditions at CRS favored the formation of polar (membrane) lipids, which have greater PUFA content. In addition, shaded conditions at CRS altered the relative proportions of ω-3 and ω-6 PUFAs in *Cladophora*, which decreased its Σω3/Σω6 ratio, but no such change was observed in any other alga or moss (Fig. 4B). The rhodophyte *Lemanea* is a common freshwater alga that in shaded and rapidly-flowing stretch of rocky streams (Sheath and Vis 2015), and which in some parts of the world constitutes an important part of people’s diet (Bhosale et al. 2012), although little is known about its FA profile. Similar to our prior studies, as well of a population of *Lemanea fluviatilis* from a stream in Turkey (Akgul et al. 2015), were high 18:1ω9 MUFA content and ω6 FAs, which confer a low Σω3/Σω6 ratio (0.5) compared to other macroalgae and periphytic diatoms (Table S1, Fig. 4, Torres-Ruiz et al. 2007). However, our data differ with regards to PUFA content. The EFAs 20:5ω3 and 20:4ω6, for example, were not detected by Akgul et al. (2015) but in our stream they constituted 6-8% and 8-10% of total FAs respectively which provide high nutritional quality (Fig. 2C). We have no other studies that have addressed local changes in the FA profile of *Lemanea*; in the shaded reach, it had significantly greater MUFA and SAFA content, a trend opposite to that of *Cladophora* and *Fontinalis* (Fig. 1C). Even though its 18:2ω6 and 18:3ω3 content remained similar independent of location, levels of 20:5ω3 and 20:4ω6 were greater in shaded conditions (Fig. 2C). Shading may have favored the production of energy-storage and membrane lipids, resulting an increase of its overall FA nutritional quality.

Overall, primary producers in the Cross River exhibited a low degree of stoichiometric homeostasis of key nutrient elements, as expected from previous studies (Frost et al. 2005). Cross River autotrophs had stoichiometric C/N and C/P ratios that were similar to lower size fractions of benthic and transported matter (Fig. 5 A, C). However, their N and P content per unit dry mass were much greater (Fig. 6). *Fontinalis dalecarlica* had the highest C/N ratio and the lowest C/P and N/P among autotrophs, making it the richest P source, especially at CRS. This bryophyte may have a greater P uptake efficiency than some species of macroalgae, a pattern observed in other bryophyte species (Pedersen et al. 2010; Pelton et al. 1998; but see Ylla et al. 2007). Even though primary producers in the Cross River had different C:N:P stoichiometric signatures, a general trend of nutritional enrichment at the shaded site was shown by N and P stoichiometry data. *Fontinalis dalecarlica*, *Cladophora* and *Lemanea* increased in P content (per unit mass) at the shaded section and *Fontinalis* and *Lemanea* also increased in N content (Figs. 5 and 6). However, their C content remained similar, which indicates the decrease in C/P and C/N ratios at CRS were due to an increase in the amount of P and N retained by these organisms. Since water column P was similar in the two reaches, local recycling processes at the shaded site may be liberating N and P into the water. Increased grazing at CRS is one possible mechanism, as macroinvertebrate egestion and excretion has been shown to result in greater periphyton N and P content (Hillebrand et al. 2004; Hillebrand et al. 2008). Another mechanism could be an increase in N and P remineralization rates by epiphytic bacteria and/or associated protozoa at the shaded section (Parry 2004; Scott et al. 2012).

### Spatial differences in quality of allochthonous matter

Spatial differences the nutritional quality of benthic organic matter varied with location and particle size, which depended in part on decomposition stage of this matter. For example, the Σω3/Σω6 ratio of UFBOM was significantly lower at the shaded reach (CRS; Fig. 4A), while CBOM had significantly greater PUFA content (e.g. 18:3ω3) in CRS (Fig. 2A, Fig. S1A). This difference in the coarse fractions may be due to difference in the sources of CBOM, which was mainly freshly-fallen green leaves in the shaded section, and decomposing leaves at the open reach. In both locations, we observed a decline in 18C PUFAs as benthic and transported matter decomposed into smaller particles, with smaller fractions having lower 18:2ω6 and 18:3ω3 content (Fig. 2A, B). In contrast, monounsaturated, saturated and bacterial FAs increased with decomposition (Fig. S1A, B). All of these patterns are consistent with previous observations of preferential degradation of PUFAs with decomposition (Torres-Ruiz and Wehr 2010). In addition, there was evidence of algal colonization of benthic organic matter in the section of the river receiving more sunlight, based on greater 20:5ω3 and higher Σω3/Σω6 ratios. These longer 20C PUFAs, especially 20:5ω3, are typical of diatoms, although the percent content was never as high as in mixed periphyton or other algal fractions (Fig. 2; Torres-Ruiz and Wehr 2020; Torres-Ruiz et al. 2007).

As expected, smaller fractions of BOM had greater stoichiometric nutritional quality with decomposition (progressively smaller fractions) at both sites, with lower C/N and C/P ratios (Fig. 5A, C). This was mainly due to a reduction of Carbon content but also to a slight increase in N and P (Fig. 6), especially in the FBOM fraction, that likely reflected microbial colonization (Farrell et al. 2018). In addition, FBOM had greater C/P and lesser P and N content at the shaded site, making it less nutritious for consumers (Figs. 5 and 6). Even though these changes were not statistically significant, they are consistent with lower EFA quality (lower algal content) observed at CRS.

Fine particulate matter drifting in the water column (FPOM and UFPOM) had greater levels of 18:2ω6 and 18:3ω3 FAs at the shaded site; this increase was not reflected in algal markers such as 20:5ω3 or Σω3/Σω6 ratio (Figs. 2B and 4A). The former are prominent in terrestrial matter (Torres-Ruiz and Wehr 2010; Torres-Ruiz et al. 2007), suggesting an allochthonous source for FPOM and UFPOM. Changes in nutritional quality of detrital organic matter was also observed in stoichiometric properties, with greater C, N and P content, higher C/N (due to disproportionately C increase) and lower C/P and N/P in the shaded location (Figs. 5 and 6). These results indicate that transported matter may have been locally enhanced by “organic matter rain” falling from the above canopy at the shaded site, either in the form of arthropods (Rozanova et al. 2019) and/or pollen (UFPOM, see Filipiak 2016). We suggest that nutrient stoichiometry was more sensitive to these changes than were FA signatures.

### Spatial differences in macroinvertebrate FA and nutrient profiles

All primary macroinvertebrate consumers in the Cross River contained algal marker FAs such as 20:5ω3, suggesting that primary producers were their main energy sources, although their relative proportions depended on species and location. Terrestrial detritus FA markers 20:0 and 22:0 concentrations were typically very low or non-detectable (0.1-0.2%; Table S2), indicating that while Carbon from terrestrial plants is being transferred through the food chain, it constituted a small part of the primary consumer diet. However, all animals contained bacterial marker FAs, with percentages ranging from 2-7%. Since bacterial markers were present in all food sources (including primary producers) it is difficult to confirm, using only FA markers, how exactly these animals are incorporating this bacterial Carbon, but in our study we suggest it most probably comes from bacterial epiphytes on primary producers.

The net-spinning caddisfly *Hydropsyche* has previously been shown to change its FA profile depending on its diet (Torres-Ruiz et al. 2010). In the Cross River, FAs in this collector reflected a diet dominated by autochthonous matter at the open site (including the moss *Fontinalis*) and a more terrestrial contribution (more SAFAs and BAFAs, less PUFAs) at the shaded site (Fig. S2A). However, a relatively high (8-12%) content of algal marker EFA 20:5ω3 at both sites suggests that autochthonous matter is an important part of its diet independent of location (Fig. 3A). Because transported matter at CRS was supplemented with omega-3 rich freshly-fallen terrestrial material, it is also likely that this matter contributed to the diet of this important collector. These patterns suggests an ability to select and assimilate food items depending on their macro (EFA) and micronutrient (P – high in *Fontinalis*) content, as well their availably. *Hydropsyche* additionally seems to be selectively incorporating the EFA 20:5ω3 as its content is higher than any of the food sources examined (Fig. 3A).

Not surprisingly, FA data show a predominantly autochthonous diet for the grazing mayfly *Epeorus*; with Σω3/Σω6 ratios that were the highest among all animals studied and all food sources available (Fig. 4C). This mayfly preferentially assimilated ω3 FAs (EFA 20:5ω3 in particular) over ω6 FAs. While maintaining a herbivorous feeding mode, its FA profile changed significantly with site, tracking the differences in the relative importance of *Cladophora* (open) and *Fontinalis* (shaded), with significantly more PUFA and less SAFA and Σω3/Σω6 ratio at the shaded site (Figs. S2A and 4). NMDS analysis placed *Epeorus* closer to *Cladophora* at the open site and to *Fontinalis* and *Lemanea* at the shaded site, likely due to differences in relative abundance of these primary producers with location. The FA data also show lower-quality mixed periphyton was not a major food source for this grazer, as shown in other systems (Mihuc and Minshall 1995). A similar pattern was seen in the grazing mayfly *Stenonema*, with high 20:5ω3 content at both sites (Figs. 1 and 4). In addition, this invertebrate tracked differences in EFA content of *Cladophora* and *Fontinalis* with site; greater PUFA and less MUFA in the shaded location (Fig. S2A). Mixed periphyton was also not major part of the diet for this grazer, similar prior studies (McWilliam-Hughes et al. 2009; Mulholland et al. 2000).

Fatty acid content of the filter feeder *Isonychia* was the least variable among primary consumers, with similar levels of all EFAs at both sites (Fig. 3A). This invertebrate had the highest levels of algal marker 20:5ω3 among all animals and the absence of a terrestrial signature with a Σω3/Σω6 ratio much greater than 1.0 (Figs. 1 and 4). Together these data suggest a preferential accumulation of these algal-based EFAs. Prior studies indicate that diatoms can be a major food item in guts of *Isonychia* (Benke 2018). Despite being classified as a filter feeder (Wallace and O’Hop 1979), its EFA-rich profile in the Cross Ross resembles that of a grazer.

*Cladophora* was the most probable food source of the case-building caddisfly *Glossosoma*, although its FA profile shows consumption of the moss *Fontinalis* as well, especially at the shaded location. *Glossosoma* had the highest amounts of specific *Cladophora* 16PUFA markers (Table S2) and was closest to this alga in the CRO NMDS plot (Fig. 1). Like other grazers, the EFA profile of this animal reflected differences in the two major producers at each site, with more PUFAs and less SAFA and MUFA at the shaded section (Fig. S2A). These results are not surprising since this caddisfly has been observed to be a grazer in other streams (McNeely et al. 2007; Mooney et al. 2016). In the Cross River, periphyton was generally of a lower nutritional quality; *Glossosoma* apparently selected other primary producers to satisfy its nutritional needs, although this species also had the highest content of bacterial FA of all animals measured (Fig. S2A), likely from epiphytic bacteria attached to *Cladophora* and *Fontinalis* (Page and Flannery 2018; Zulkifly et al. 2012). Our data do not support the general conclusion based on stable isotope tracer studies that the source of bacterial Carbon within food webs is usually bacteria colonizing terrestrial detritus (Hall 1995; Sánchez-Carrillo and Álvarez-Cobelas 2018).

The water penny *Psephenus* also had relatively high content of 16PUFA *Cladophora* markers (Table S2), and reflected changes in FA content similar to this alga and *Fontinalis*. However, and even though it was still greater than 1.0, its Σω3/Σω6 ratio was the lowest among all primary consumers (Fig. 4 C). This ratio has been proposed as the marker of relative consumption of autochthonous matter rich in omega-3 FAs vs. microbially colonized terrestrial matter rich in omega-6 FAs (Torres-Ruiz and Wehr 2010). Results from the present study challenge this paradigm. Several primary producers, in particular *Fontinalis* and *Lemanea,* which are rich in omega-6 FAs, especially the essential 20:4ω6, have Σω3/Σω6 ratios < 1.0 (Fig. 4 B). It is clear that some consumers preferentially accumulate omega-3 FAs (especially 20:5ω3) and as a result have very high Σω3/Σω6 ratios, but we suggest that a lower Σω3/Σω6 ratio, as in the case of *Psephenus*, may not mean a reliance on detrital sources, but a dependence on different primary producers, such as the bryophytes or red algae.

In this study, we have considered the insects *Rhyacophila, Paragnetina* and *Nigronia* predators, based on extensive prior research (Fuller and Hynes 1987; Manuel and Folsom 1982; Tavares-Cromar and Williams 1996). FA data alone were not capable of discerning trophic position, however, it is clear that these predators fed on primary consumers, as they plotted near them in the NMDS plots for both CRO and CRS (Fig. 1), with similar high content of PUFAs, and lower SAFAs and bacterial FAs (Fig. S2B). There were subtle differences present in the FA profile of the megalopteran *Nigronia* with greater 18:2ω6 and 20:4ω6, and lower 18:3ω3 than the other predators, which resulted in an Σω3/Σω6 ratio closer to 1.0 (Figs. 3B and 4C). Despite these differences, it is clear that predators were relying on insects that had consumed primary producers *Cladophora* and *Fontinalis*. It is also evident from their FA profile that a small portion of their FAs came from detrital matter (20:0 and 22:0, 0.1-0.9%, Table S2); these predators may consume small numbers of detritivores (e.g. oligochaetes isopods), a group were did not analyze here.

Exposure to sunlight or shade apparently had smaller effects on predator nutrition. Only the stonefly *Paragnetina* had significant greater PUFAs (18:3ω3, 20:4ω6 and 20:5ω3, Fig. 3B), and less SAFA (Fig. S2B) at the shaded location. In contrast, *Nigronia* and *Rhyacophila* had similar FA profiles at both locations (Figs. 3B and S2B). We assume that these predators regulate their FA profile to a greater extent than primary producers by assimilating and/or synthesizing the essential FAs they need, irrespective of their diet they consume, as long as it contains the FA building blocks needed for synthesis (EFAs such as 18:2ω6, 318:3ω3, 20:4ω6 and 20:5ω3, present in all primary consumers).

With respect to their nutrient ratios, most invertebrates were relatively homeostatic, with C/N and C/P ratios similar in both stream reaches and similar to those observed in prior studies (Bowman et al. 2005; Cross et al. 2003; Lauridsen et al. 2012; Ohta et al. 2016). These ratios reveal a strong N and P limitation with respect to most terrestrially-derived food sources (Fig. 5) and a lower imbalance with respect to smaller fractions of detritus and autochthonous matter. The ratios of most allochthonous matter in the present study were considerably greater published thresholds for elemental ratios required for aquatic consumers (Frost et al. 2006; Halvorson et al. 2015). However, in terms of nutrient content per unit mass, autochthonous sources were much closer to invertebrates than any fraction of detrital matter (Fig. 6). This contradiction requires that consumers of fine or ultrafine benthic organic matter would need to consume large quantities to meet nutritional needs. Interestingly, the moss *Fontinalis* had the closest C/P ratio (and highest P content) to invertebrates. Perhaps it is not surprising that FA markers indicate bryophytes as a common food source for most of the invertebrates in the present study (Kalachova et al. 2011; McWilliam-Hughes et al. 2009; Torres-Ruiz and Wehr 2020). As in our previous study (Torres-Ruiz and Wehr 2020), our data here indicate that that many of not most animals depend to an important degree on autochthonous food items, which provide not only elemental nutrients, but also essential macromolecules needed for their growth and reproduction. However, N and P are still limiting in this food web. Interestingly, some animals in our study had C/P ratios that were 50-100% higher than the rest, particularly at CRS (mainly *Psephenus* and *Rhyacophila*). This is may be due to differences in size and/or growth rate (Cross et al. 2005; Demi et al. 2018; Sterner and Elser 2002). In addition, both of these animals were non-homeostatic with respect to their C/P and N/P ratios, with greater P content at the shaded site, in which autotrophs also had greater P content (Figs. 5 and 6). This suggests many stream invertebrates may be able to adjust their P content depending on the P content of their diet, and thereby minimize nutrient limitation (Demi et al. 2018; Small and Pringle 2010).

This study further confirmed the importance of autochthonous food sources for low-order stream invertebrates, and has highlighted the importance of processes driven by local differences in light availability and canopy-derived nutrient-rich matter in stream food webs. These small-scale differences are tracked differently by each consumer, patterns that also depend on the nutrient examined. Invertebrates exhibit a great degree of homeostasis with respect to N and P, but were more varied with regard to essential FAs, which also depended on trophic level.

## References

1. Akgul, R., B. Kizilkaya, F. Akgul & H. Erdugan, 2015. Total lipid and fatty acid composition of twelve algae from Çanakkale (Turkey). Indian Journal of Geo-Marine Science 44(4):495–500 doi:http://nopr.niscair.res.in/handle/123456789/34725.

2. Benke, A. C., 2018. River food webs: an integrative approach to bottom-up flow webs, top-down impact webs, and trophic position. Ecology 99(6):1370–1381 doi:http://doi.org/10.1002/ecy.2228.

3. Berg, M. P. & J. Bengtsson, 2007. Temporal and spatial variability in soil food web structure. Oikos 116(11):1789–1804 doi:http://doi.org/10.1111/j.0030-1299.2007.15748.x.

4. Bhosale, R. A., J. Rout & B. B. Chaugule, 2012. The Ethnobotanical study of an edible freshwater red alga, *Lemanea fluviatilis* (L.) C. Ag. from Manipur, India. Ethnobotany Research and Applications 10:69–76 doi:http://doi.org/10.17348/ERA.10.0.069-076.

5. Bing, T., J. Müller, B. Glaser, R. Brandl & M. Brändle, 2015. Variation in diet across an elevational gradient in the larvae of two *Hydropsyche* species (Trichoptera). Limnologica 52:83–88 doi:https://doi.org/10.1016/j.limno.2015.04.001.

6. Bowman, M. F., P. A. Chambers & D. W. Schindler, 2005. Changes in stoichiometric constraints on epilithon and benthic macroinvertebrates in response to slight nutrient enrichment of mountain rivers. Freshwater Biology 50(11):1836–1852 doi:https://doi.org/10.1111/j.1365-2427.2005.01454.x.

7. Cashman, M. J., F. Pilotto, G. L. Harvey, G. Wharton & M. T. Pusch, 2016. Combined stable-isotope and fatty-acid analyses demonstrate that large wood increases the autochthonous trophic base of a macroinvertebrate assemblage. Freshwater biology 61(4):549–564 doi:https://doi.org/10.1111/fwb.12727.

8. Cashman, M. J., J. D. Wehr & K. Truhn, 2013. Elevated light and nutrients alter the nutritional quality of stream periphyton. Freshwater Biology 58(7):1447–1457 doi: https://doi.org/10.1111/fwb.12142.

9. Closs, G. & P. Lake, 1994. Spatial and temporal variation in the structure of an intermittent-stream food web. Ecological monographs 64(1):1–21 doi:https://doi.org/10.2307/2937053.

10. Cross, W., B. Johnson, J. Wallace & A. Rosemond, 2005. Contrasting response of stream detritivores to long-term nutrient enrichment. Limnology and Oceanography 50(6):1730–1739 doi: https://doi.org/10.4319/lo.2005.50.6.1730.

11. Cross, W. F., J. P. Benstead, A. D. Rosemond & J. Bruce Wallace, 2003. Consumer-resource stoichiometry in detritus-based streams. Ecology Letters 6(8):721–732 doi:https://doi.org/10.1046/j.1461-0248.2003.00481.x.

12. Demi, L. M., J. P. Benstead, A. D. Rosemond & J. C. Maerz, 2018. Litter P content drives consumer production in detritus-based streams spanning an experimental N:P gradient. Ecology 99(2):347–359 doi:http://doi.org/10.1002/ecy.2118.

13. Desvilettes, C., G. Bourdier, J. C. Breton & P. Combrouze, 1994. Fatty acids as organic markers for the study of trophic relationships in littoral cladoceran communities of a pond. Journal of Plankton Research 16(6):643–659 doi:https://doi.org/10.1093/plankt/16.6.643.

14. Dickman, E. M., M. J. Vanni & M. J. Horgan, 2006. Interactive effects of light and nutrients on phytoplankton stoichiometry. Oecologia 149(4):676–689 doi:http://doi.org/10.1007/s00442-006-0473-5.

15. Dodds, W. K., S. A. Higgs, M. J. Spangler, J. Guinnip, J. D. Scott, S. C. Hedden, B. D. Frenette, R. Taylor, A. E. Schechner, D. J. Hoeinghaus & M. A. Evans-White, 2018. Spatial heterogeneity and controls of ecosystem metabolism in a Great Plains river network. Hydrobiologia 813(1):85–102 doi:http://doi.org/10.1007/s10750-018-3516-0.

16. Filipiak, M., 2016. Pollen stoichiometry may influence detrital terrestrial and aquatic food webs. Frontiers in Ecology and Evolution 4:1–8.

17. Findlay, S., M. Pace & D. Fischer, 1996. Spatial and temporal variability in the lower food web of the tidal freshwater Hudson River. Estuaries 19(4):866–873 doi:https://doi.org/10.2307/1352303.

18. Finlay, J. C., S. Khandwala & M. E. Power, 2002. Spatial scales of carbon flow in a river food web. Ecology 83(7):1845–1859 doi:https://doi.org/10.1890/0012-9658(2002)083[1845:SSOCFI]2.0.CO;2.

19. Franken, R. J., B. Waluto, E. T. Peeters, J. J. Gardeniers, J. A. Beijer & M. Scheffer, 2005. Growth of shredders on leaf litter biofilms: the effect of light intensity. Freshwater Biology 50(3):459–466 doi:https://doi.org/10.1111/j.1365-2427.2005.01333.x.

20. Frost, P. C., J. P. Benstead, W. F. Cross, H. Hillebrand, J. H. Larson, M. A. Xenopoulos & T. Yoshida, 2006. Threshold elemental ratios of Carbon and Phosphorus in aquatic consumers. Ecology Letters 9(7):774–779 doi:http://doi.org/10.1111/j.1461-0248.2006.00919.x.

21. Frost, P. C., W. F. Cross & J. P. Benstead, 2005. Ecological stoichiometry in freshwater benthic ecosystems: an introduction. Freshwater Biology 50(11):1781–1785.

22. Fuller, R. L. & H. Hynes, 1987. Feeding ecology of three predacious aquatic insects and two fish in a riffle of the Speed River, Ontario. Hydrobiologia 150(3):243–255 doi:http://doi.org/10.1007/BF00008706.

23. Gulis, V., K. A. Kuehn, L. N. Schoettle, D. Leach, J. P. Benstead & A. D. Rosemond, 2017. Changes in nutrient stoichiometry, elemental homeostasis and growth rate of aquatic litter-associated fungi in response to inorganic nutrient supply. The ISME Journal 11(12):2729–2739 doi:http://doi.org/10.1038/ismej.2017.123.

24. Guo, F., S. E. Bunn, M. T. Brett, B. Fry, H. Hager, X. Ouyang & M. J. Kainz, 2018. Feeding strategies for the acquisition of high-quality food sources in stream macroinvertebrates: Collecting, integrating, and mixed feeding. Limnology and Oceanography 63(5):1964–1978 doi:http://doi.org/10.1002/lno.10818.

25. Guo, F., S. E. Bunn, M. T. Brett & M. J. Kainz, 2017. Polyunsaturated fatty acids in stream food webs–high dissimilarity among producers and consumers. Freshwater Biology 62(8):1325–1334 doi: https://doi.org/10.1111/fwb.12956.

26. Guo, F., M. J. Kainz, F. Sheldon & S. E. Bunn, 2016a. Effects of light and nutrients on periphyton and the fatty acid composition and somatic growth of invertebrate grazers in subtropical streams. Oecologia 181(2):449–462 doi:http://doi.org/10.1007/s00442-016-3573-x.

27. Guo, F., M. J. Kainz, F. Sheldon & S. E. Bunn, 2016b. The importance of high-quality algal food sources in stream food webs–current status and future perspectives. Freshwater Biology 61(6):815–831 doi:https://doi.org/10.1111/fwb.12755.

28. Guo, F., M. J. Kainz, D. Valdez, F. Sheldon & S. E. Bunn, 2016c. The effect of light and nutrients on algal food quality and their consequent effect on grazer growth in subtropical streams. Freshwater Science 35(4):1202–1212.

29. Guo, F., M. J. Kainz, D. Valdez, F. Sheldon & S. E. Bunn, 2016d. High-quality algae attached to leaf litter boost invertebrate shredder growth. Freshwater Science 35(4):1213–1221 doi:http://doi.org/10.1086/688667.

30. Hall, R. O., 1995. Use of a stable carbon isotope addition to trace bacterial carbon through a stream food web. Journal of the North American Benthological Society 14(2):269–277 doi:http://doi.org/10.2307/1467779.

31. Halvorson, H. M., C. L. Fuller, S. A. Entrekin, J. T. Scott & M. A. Evans-White, 2019. Interspecific homeostatic regulation and growth across aquatic invertebrate detritivores: a test of ecological stoichiometry theory. Oecologia 190(1):229–242 doi:http://doi.org10.1007/s00442-019-04409-w.

32. Halvorson, H. M., J. T. Scott, A. J. Sanders & M. A. Evans-White, 2015. A stream insect detritivore violates common assumptions of threshold elemental ratio bioenergetics models. Freshwater Science 34(2):508–518 doi:https://doi.org/10.1086/680724.

33. Harlıoğlu, M. M., K. Köprücü, A. G. Harlıoğlu, Ö. Yılmaz, S. Mişe Yonar, S. Aydın & T. Çakmak Duran, 2015. Effects of dietary n-3 polyunsaturated fatty acids on the nutritional quality of abdomen meat and hepatopancreas in a freshwater crayfish (*Astacus leptodactylu*). Journal of Food Composition and Analysis 41:144–150 doi:http://doi.org/10.1016/j.jfca.2015.01.011.

34. Hill, W. R., J. Rinchard & S. Czesny, 2011. Light, nutrients and the fatty acid composition of stream periphyton. Freshwater Biology 56(9):1825–1836 doi:https://doi.org/10.1111/j.1365-2427.2011.02622.x.

35. Hill, W. R., M. G. Ryon & E. M. Schilling, 1995. Light limitation in a stream ecosystem: responses by primary producers and consumers. Ecology 76(4):1297–1309 doi:https://doi.org/10.2307/1940936.

36. Hillebrand, H., G. De Montpellier & A. Liess, 2004. Effects of macrograzers and light on periphyton stoichiometry. Oikos 106(1):93–104 doi:https://doi.org/10.1111/j.0030-1299.2004.13166.x.

37. Hillebrand, H., P. Frost & A. Liess, 2008. Ecological stoichiometry of indirect grazer effects on periphyton nutrient content. Oecologia 155(3):619–630 doi:http://doi.org/10.1007/s00442-007-0930-9.

38. Hu, Q., M. Sommerfeld, E. Jarvis, M. Ghirardi, M. Posewitz, M. Seibert & A. Darzins, 2008. Microalgal triacylglycerols as feedstocks for biofuel production: perspectives and advances. The Plant Journal 54(4):621–639 doi:http://doi.org/10.1111/j.1365-313X.2008.03492.x.

39. Huggins, K., J. J. Frenette & M. T. Arts, 2004. Nutritional quality of biofilms with respect to light regime in Lake Saint-Pierre (Québec, Canada). Freshwater Biology 49(7):945–959 doi: https://doi.org/10.1111/j.1365-2427.2004.01236.x.

40. Jamieson, G. R. & E. H. Reid, 1976. Lipids of *Fontinalis antipyretica*. Phytochemistry 15(11):1731–1734 doi:https://doi.org/10.1016/S0031-9422(00)97466-1.

41. Kalacheva, G., N. Sushchik, M. Gladyshev & O. Makhutova, 2009. Seasonal dynamics of fatty acids in the lipids of water moss *Fontinalis antipyretica* from the Yenisei River. Russian Journal of Plant Physiology 56(6):795–807 doi:https://doi.org/10.1134/S1021443709060090.

42. Kalachova, G., M. Gladyshev, N. Sushchik & O. Makhutova, 2011. Water moss as a food item of the zoobenthos in the Yenisei River. Open Life Sciences 6(2):236–245 doi:https://doi.org/10.2478/s11535-010-0115-0.

43. Kharlamenko, V. I., N. V. Zhukova, S. V. Khotimchenko, V. I. Svetashev & G. M. Kamenev, 1995. Fatty acids as markers of food sources in a shallow-water hydrothermal ecosystem (Kraternaya Bight, Yankich Island, Kurile Islands). Marine Ecology Progress Series 120(1-3):231–241 doi: http://doi.org/10.3354/meps120231

44. Lancaster, J., D. C. Bradley, A. Hogan & S. Waldron, 2005. Intraguild omnivory in predatory stream insects. Journal of Animal Ecology 74(4):619–629.

45. Lauridsen, R. B., F. K. Edwards, M. J. Bowes, G. Woodward, A. G. Hildrew, A. T. Ibbotson & J. I. Jones, 2012. Consumer–resource elemental imbalances in a nutrient-rich stream. Freshwater Science 31(2):408–422 doi:https://doi.org/10.1899/11-052.

46. Liess, A. & H. Hillebrand, 2006. Role of nutrient supply in grazer–periphyton interactions: reciprocal influences of periphyton and grazer nutrient stoichiometry. Journal of the North American Benthological Society 25(3):632–642 doi:https://doi.org/10.1899/0887-3593(2006)25[632:RONSIG]2.0.CO;2.

47. Liu, J., P. Vanormelingen & W. Vyverman, 2016. Fatty acid profiles of four filamentous green algae under varying culture conditions. Bioresource Technology 200:1080–1084 doi:http://doi.org/10.1016/j.biortech.2015.11.001.

48. Manfrin, A., L. Traversetti, F. Pilotto, S. Larsen & M. Scalici, 2016. Effect of spatial scale on macroinvertebrate assemblages along a Mediterranean river. Hydrobiologia 765(1):185–196 doi:https://doi.org/10.1007/s10750-015-2412-0.

49. Manuel, K. L. & T. C. Folsom, 1982. Instar sizes, life cycles, and food habits of five *Rhyacophila* (Trichoptera: Rhyacophilidae) species from the Appalachian Mountains of South Carolina, U.S.A. Hydrobiologia 97(3):281–285 doi:http://doi.org/10.1007/bf00007115.

50. Martyniuk, N., B. Modenutti & E. G. Balseiro, 2019. Light intensity regulates stoichiometry of benthic grazers through changes in the quality of stream periphyton. Freshwater Science 38(2):391–405.

51. McNeely, C., J. C. Finlay & M. E. Power, 2007. Grazer traits, competition, and carbon sources to a headwater-stream food web. Ecology 88(2):391–401 doi:http://doi.org/10.1890/0012-9658(2007)88[391:gtcacs]2.0.co;2.

52. McWilliam-Hughes, S. M., T. D. Jardine & R. A. Cunjak, 2009. Instream C sources for primary consumers in two temperate, oligotrophic rivers: possible evidence of bryophytes as a food source. Journal of the North American Benthological Society 28(3):733–743 doi:https://doi.org/10.1899/08-103.1.

53. Mihuc, T. B. & G. W. Minshall, 1995. Trophic Generalists vs. Trophic Specialists: Implications for Food Web Dynamics in Post-Fire Streams. Ecology 76(8):2361–2372 doi:http://doi.org/10.2307/2265813.

54. Mille-Lindblom, C., H. Fischer & L. J. Tranvik, 2006. Antagonism between bacteria and fungi: substrate competition and a possible tradeoff between fungal growth and tolerance towards bacteria. Oikos 113(2):233–242 doi:https://doi.org/10.1111/j.2006.0030-1299.14337.x.

55. Miura, A. & J. Urabe, 2015. Spatial and seasonal changes in species diversity of epilithic fungi along environmental gradients of a river. Freshwater Biology 60(4):673–685 doi:https://doi.org/10.1111/fwb.12514.

56. Mooney, R. J., E. A. Strauss & R. J. Haro, 2016. Nutrient-specific foraging by *Glossosoma intermedium* larvae leads to conspecific case grazing. Freshwater Science 35(3):873–881 doi:https://doi.org/10.1086/686699.

57. Mulholland, P. J., J. L. Tank, D. M. Sanzone, W. M. Wollheim, B. J. Peterson, J. R. Webster & J. L. Meyer, 2000. Food resources of stream macroinvertebrates determined by natural-abundance stable C and N isotopes and a ^15^N tracer addition. Journal of the North American Benthological Society 19(1):145–157 doi:https://doi.org/10.2307/1468287.

58. Napolitano, G. E., 1994. The relationship of lipids with light and chlorophyll measurements in freshwater algae and periphyton. Journal of Phycology 30(6):943–950 doi:https://doi.org/10.1111/j.0022-3646.1994.00943.x.

59. Napolitano, G. E., R. J. Pollero, A. M. Gayoso, B. A. MacDonald & R. J. Thompson, 1997. Fatty acids as trophic markers of phytoplankton blooms in the Bahia Blanca estuary (Buenos Aires, Argentina) and in Trinity Bay (Newfoundland, Canada). Biochemical Systematics and Ecology 25(8):739–755 doi:https://doi.org/10.1016/S0305-1978(97)00053-7.

60. Ohta, T., S. Matsunaga, S. Niwa, K. Kawamura & T. Hiura, 2016. Detritivore stoichiometric diversity alters litter processing efficiency in a freshwater ecosystem. Oikos 125(8):1162–1172 doi:https://doi.org/10.1111/oik.02788.

61. Page, K. A. & M. K. Flannery, 2018. Microbial Epiphytes of Deep-Water Moss in Crater Lake, Oregon. Northwest Science 92(4):240–250 doi:https://doi.org/10.3955/046.092.0402.

62. Parry, J. D., 2004. Protozoan grazing of freshwater biofilms. In Laskin, A. I., J. W. Bennett & G. M. Gadd (eds) Advances in applied microbiology. vol 54. Elsevier Academic Press, California, 167–196.

63. Pedersen, M. F., J. Borum & F. L. Fotel, 2010. Phosphorus dynamics and limitation of fast-and slow-growing temperate seaweeds in Oslofjord, Norway. Marine Ecology Progress Series 399:103–115 doi:https://doi.org/10.3354/meps08350.

64. Pelton, D. K., S. N. Levine & M. Braner, 1998. Measurements of Phosphorus uptake by macrophytes and epiphytes from the LaPlatte River (VT) using 32P in stream microcosms. Freshwater Biology 39(2):285–299 doi:https://doi.org/10.1046/j.1365-2427.1998.00281.x.

65. Persson, J., P. Fink, A. Goto, J. M. Hood, J. Jonas & S. Kato, 2010. To be or not to be what you eat: regulation of stoichiometric homeostasis among autotrophs and heterotrophs. Oikos 119(5):741–751 doi:https://doi.org/10.1111/j.1600-0706.2009.18545.x.

66. Rozanova, O. L., S. M. Tsurikov, A. V. Tiunov & E. E. Semenina, 2019. Arthropod rain in a temperate forest: Intensity and composition. Pedobiologia 75:52–56.

67. Sabater, S., J. Artigas, A. Gaudes, I. Munoz, G. Urrea & A. M. Romani, 2011. Long-term moderate nutrient inputs enhance autotrophy in a forested Mediterranean stream. Freshwater Biology 56(7):1266–1280 doi:https://doi.org/10.1111/j.1365-2427.2010.02567.x.

68. Sanches, L. F., R. D. Guariento, A. Caliman, R. L. Bozelli & F. A. Esteves, 2011. Effects of nutrients and light on periphytic biomass and nutrient stoichiometry in a tropical black-water aquatic ecosystem. Hydrobiologia 669(1):35–44.

69. Sánchez-Carrillo, S. & M. Álvarez-Cobelas, 2018. Stable isotopes as tracers in aquatic ecosystems. Environmental Reviews 26(1):69–81 doi:http://dx.doi.org/10.1139/er-2017-0040.

70. Schagerl, M. & M. Kerschbaumer, 2009. Autecology and morphology of selected Vaucheria species (Xanthophyceae). Aquatic Ecology 43(2):295–303 doi:https://doi.org/10.1007/s10452-007-9163-6.

71. Scott, J. T., J. B. Cotner & T. M. Lapara, 2012. Variable stoichiometry and homeostatic regulation of bacterial biomass elemental composition. Frontiers in Microbiology 3:1–8 doi:http://doi.org/10.3389/fmicb.2012.00042.

72. Sheath, R. G. & M. L. Vis, 2015. Red Algae. In Wehr, J. D., R. G. Sheath & J. P. Kociolek (eds) Freshwater Algae of North America (Second Edition). Academic Press, Boston, 237–264.

73. Small, G. E. & C. M. Pringle, 2010. Deviation from strict homeostasis across multiple trophic levels in an invertebrate consumer assemblage exposed to high chronic phosphorus enrichment in a Neotropical stream. Oecologia 162(3):581–590 doi:http://doi.org/10.1007/s00442-009-1489-4.

74. Sokal, R. R. & F. Rohlf, 1995. Biometry: the principles and practice of statistics in biological research. Freeman and Company, New York.

75. Solórzano, L. & J. H. Sharp, 1980. Determination of total dissolved Phosphorus and particulate Phosphorus in natural waters. Limnology and Oceanography 25(4):754–758 doi:https://doi.org/10.4319/lo.1980.25.4.0754.

76. Steinman, A. D. & C. D. McIntire, 1987. Effects of irradiance on the community structure and biomass of algal assemblages in laboratory streams. Canadian journal of fisheries and aquatic sciences 44(9):1640–1648 doi: https://doi.org/10.1139/f87-199.

77. Sterner, R. W. & J. J. Elser, 2002. Ecological Stoichiometry. The biology of elements from molecules to the biosphere, 1 edn. Princeton University Press, Princeton, NJ.

78. Sterner, R. W., J. J. Elser, E. J. Fee, S. J. Guildford & T. H. Chrzanowski, 1997. The light: nutrient ratio in lakes: the balance of energy and materials affects ecosystem structure and process. The American Naturalist 150(6):663–684 doi:http://doi.org/10.1086/286088.

79. Sushchik, N. N., M. I. Gladyshev, A. V. Moskvichova, O. N. Makhutova & G. S. Kalachova, 2003. Comparison of fatty acid composition in major lipid classes of the dominant benthic invertebrates of the Yenisei river. Comparative biochemistry and physiology Part B, Biochemistry & molecular biology 134(1):111–122 doi:http://doi.org/10.1016/s1096-4959(02)00191-4.

80. Taube, R., L. Ganzert, H.-P. Grossart, G. Gleixner & K. Premke, 2018. Organic matter quality structures benthic fatty acid patterns and the abundance of fungi and bacteria in temperate lakes. Science of the Total Environment 610:469–481 doi:https://doi.org/10.1016/j.scitotenv.2017.07.256.

81. Tavares-Cromar, A. F. & D. D. Williams, 1996. The importance of temporal resolution in food web analysis: evidence from a detritus-based stream. Ecological Monographs 66(1):91–113 doi:https://doi.org/10.2307/2963482.

82. Torres-Ruiz, M. & J. D. Wehr, 2010. Changes in the nutritional quality of decaying leaf litter in a stream based on fatty acid content. Hydrobiologia 651(1):265–278 doi:http://doi.org/10.1007/s10750-010-0305-9.

83. Torres-Ruiz, M. & J. D. Wehr, 2020. Complementary information from fatty acid and nutrient stoichiometry data improve stream food web analyses. Hydrobiologia 847(2):629–645 doi:https://doi.org/10.1007/s10750-019-04126-8.

84. Torres-Ruiz, M., J. D. Wehr & A. A. Perrone, 2007. Trophic relations in a stream food web: importance of fatty acids for macroinvertebrate consumers. Journal of the North American Benthological Society 26(3):509–522 doi:https://doi.org/10.1899/06-070.1.

85. Torres-Ruiz, M., J. D. Wehr & A. A. Perrone, 2010. Are net-spinning caddisflies what they eat? An investigation using controlled diets and fatty acids. Journal of the North American Benthological Society 29(3):803–813 doi:https://doi.org/10.1899/09-162.1.

86. Tsoi, W. Y., W. L. Hadwen & C. S. Fellows, 2011. Spatial and temporal variation in the ecological stoichiometry of aquatic organisms in an urban catchment. Journal of the North American Benthological Society 30(2):533–545 doi:https://doi.org/10.1899/10-085.1.

87. Twining, C. W., J. T. Brenna, N. G. Hairston Jr & A. S. Flecker, 2016. Highly unsaturated fatty acids in nature: what we know and what we need to learn. Oikos 125(6):749–760 doi:https://doi.org/10.1111/oik.02910.

88. Twining, C. W., D. C. Josephson, C. E. Kraft, J. T. Brenna, P. Lawrence & A. S. Flecker, 2017. Limited seasonal variation in food quality and foodweb structure in an Adirondack stream: insights from fatty acids. Freshwater Science 36(4):877–892 doi:https://doi.org/10.1086/694335.

89. Volk, C. & P. Kiffney, 2012. Comparison of fatty acids and elemental nutrients in periphyton, invertebrates, and cutthroat trout (*Oncorhynchus clarki*) in conifer and alder streams of western Washington state. Aquatic Ecology 46(1):85–99 doi:http://doi.org/10.1007/s10452-011-9383-7.

90. Wallace, J. B. & J. O’Hop, 1979. Fine particle suspension-feeding capabilities of *Isonychia* spp.(Ephemeroptera: Siphlonuridae). Annals of the entomological Society of America 72(3):353–357 doi:https://doi.org/10.1093/aesa/72.3.353.

91. Wehr, J. D., A. Empain, C. Mouvet, P. J. Say & B. A. Whitton, 1983. Methods for processing aquatic mosses used as monitors of heavy metals. Water Research 17(9):985–992 doi:https://doi.org/10.1016/0043-1354(83)90038-6.

92. Wellnitz, T. & R. B. Rader, 2003. Mechanisms influencing community composition and succession in mountain stream periphyton: interactions between scouring history, grazing, and irradiance. Journal of the North American Benthological Society 22(4):528–541 doi:https://doi.org/10.2307/1468350.

93. Wetzel, R., 2005. Periphyton in the aquatic ecosystem and food webs. In Azim, M. E. V., M. C. J.; van Dam, A. A.; Beveridge, M. C. M (ed) Periphyton: ecology, exploitation, and management CABI Publishing, London, 51–69.

94. Winemiller, K. O., A. S. Flecker & D. J. Hoeinghaus, 2010. Patch dynamics and environmental heterogeneity in lotic ecosystems. Journal of the North American Benthological Society 29(1):84–99 doi:https://doi.org/10.1899/08-048.1.

95. Wyatt, K. H., R. C. Seballos, M. N. Shoemaker, S. P. Brown, S. Chandra, K. A. Kuehn, A. R. Rober & S. Sadro, 2019. Resource constraints highlight complex microbial interactions during lake biofilm development. Journal of Ecology 107(6):2737–2746 doi:https://doi.org/10.1111/1365-2745.13223.

96. Ylla, I., A. M. Romaní & S. Sabater, 2007. Differential effects of nutrients and light on the primary production of stream algae and mosses. Fundamental and Applied Limnology/Archiv für Hydrobiologie 170(1):1–10 doi:http://doi.org/10.1127/1863-9135/2007/0170-001.

97. Zhang, P., R. F. van den Berg, C. H. van Leeuwen, B. A. Blonk & E. S. Bakker, 2018. Aquatic omnivores shift their trophic position towards increased plant consumption as plant stoichiometry becomes more similar to their body stoichiometry. PloS one 13(9):e0204116 doi:https://doi.org/10.1371/journal.pone.0204116.

98. Zulkifly, S., A. Hanshew, E. B. Young, P. Lee, M. E. Graham, M. E. Graham, M. Piotrowski & L. E. Graham, 2012. The epiphytic microbiota of the globally widespread macroalga *Cladophora glomerata* (Chlorophyta, Cladophorales). American Journal of Botany 99(9):1541–1552 doi:http://doi.org/10.3732/ajb.1200161.

