## Supplementary Information for "Spatial differences in elemental stoichiometry and essential fatty acid content of food sources and consumers in a stream food web"

**Index**

- Supplementary Figure 1
- Supplementary Figure 2
- Supplementary Table 1
- Supplementary Table 2

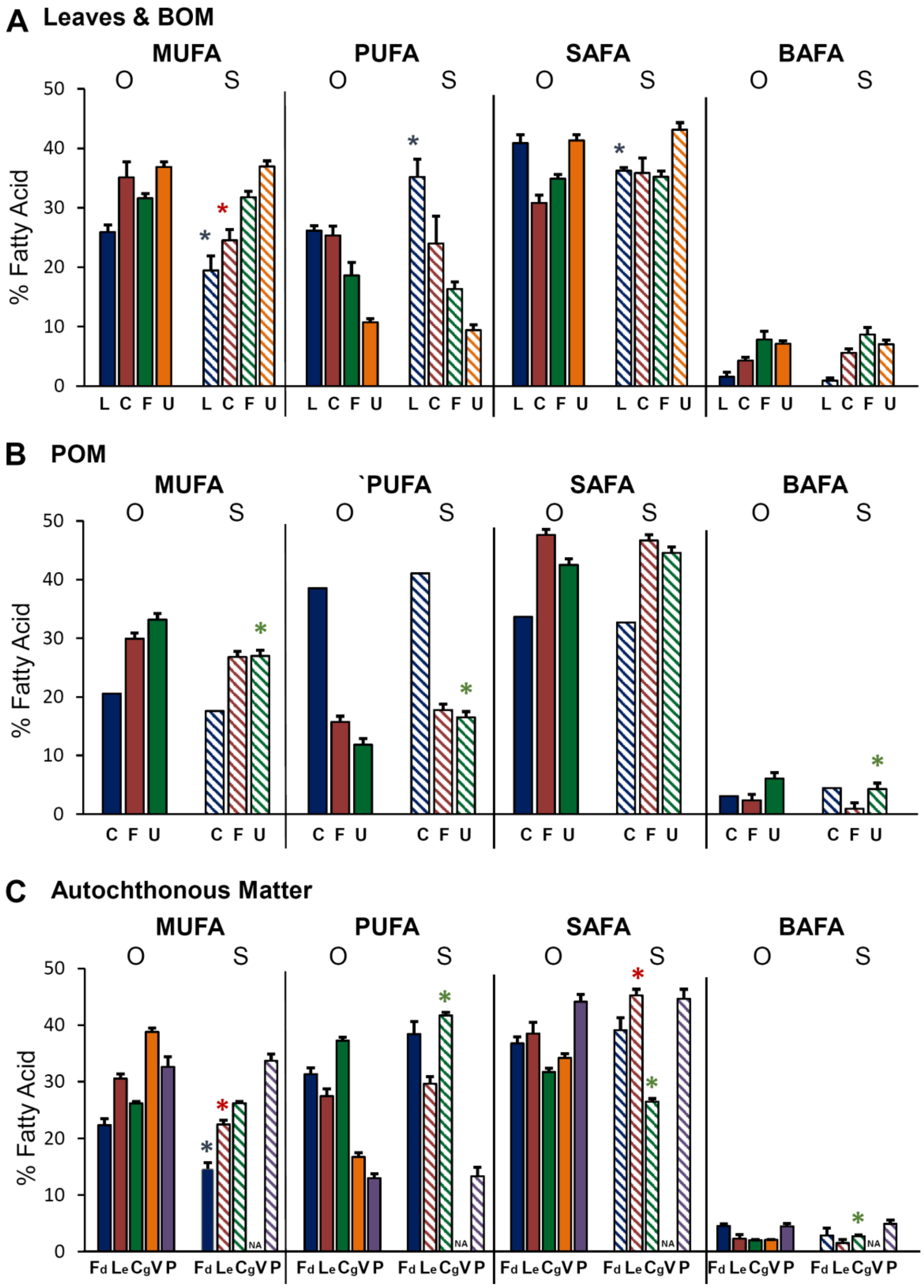


**Figure S1.** Changes with site of mono unsaturated fatty acids (MUFA), polyunsaturated fatty acids (PUFA), saturated fatty acids (SAFA) and bacterial fatty acids (BAFA) content of leaves and benthic matter **(A),** particulate matter **(B)** and autochthonous matter **(C)** in the open (O; solid bars) and shaded (S; striped bars) areas of the Cross River. Values represent means (±1 SE). Asterisk above bars represent significant differences with site ($p < 0.05$). L = leaves; C = coarse; F = fine; U = ultrafine; Fd = *Fontinalis*; Le = *Lemanea*; Cg = *Cladophora*; V = *Vaucheria*; P = periphyton.

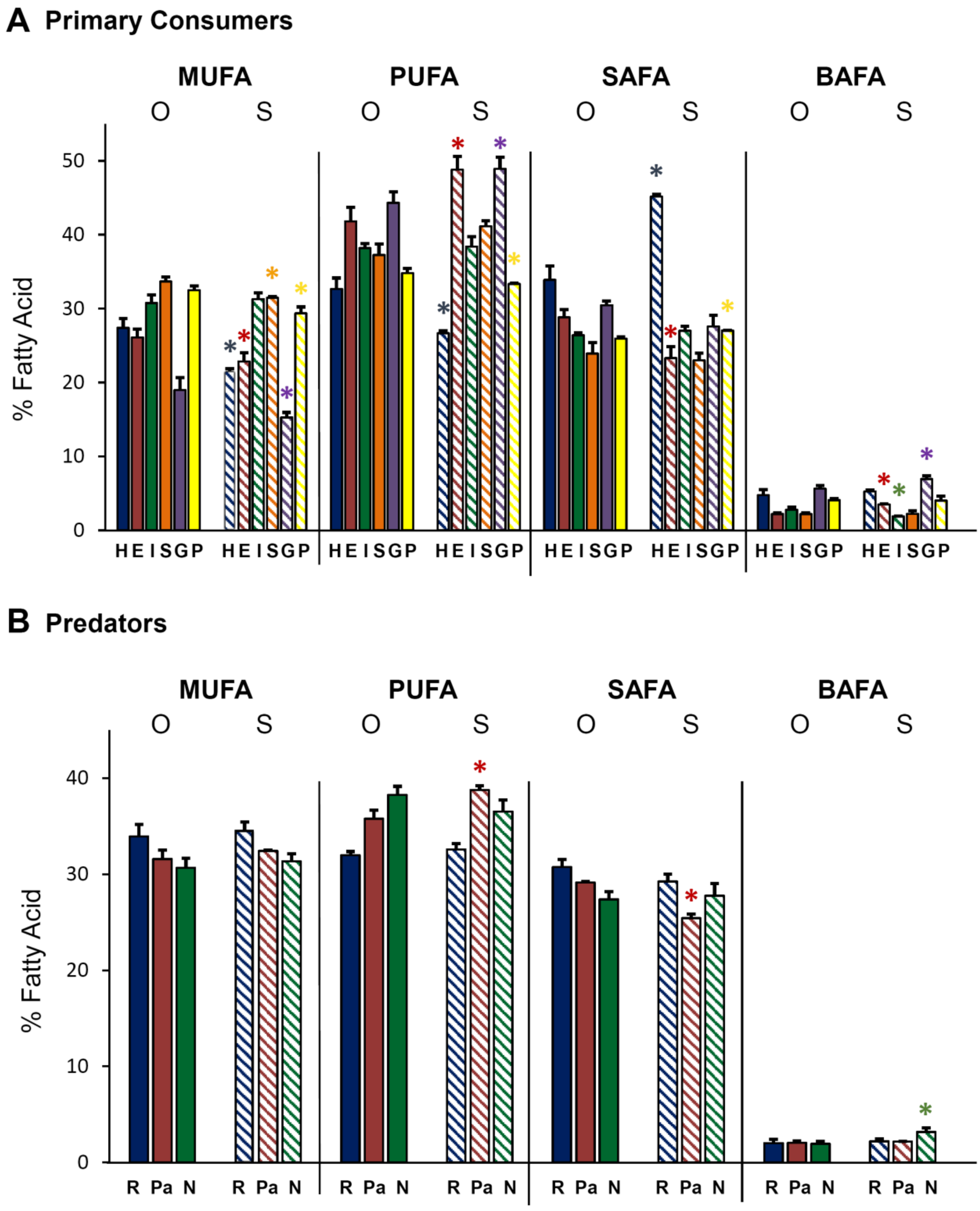


**Figure S2.** Changes with site of mono unsaturated fatty acids (MUFA), polyunsaturated fatty acids (PUFA), saturated fatty acids (SAFA) and bacterial fatty acids (BAFA) content of primary consumers **(A)** and predators **(B)** in the open (O; solid bars) and shaded (S; striped bars) areas of the Cross River. Values represent means (±1 SE). Asterisk above bars represent significant differences with site ($p < 0.05$). H = *Hydropsyche*; E = *Epeorus*; I = *Isonychia*; S = *Stenonema*; G = *Glossosoma*; P = *Psephenus*; R = *Rhyacophila*; Pa = *Paragnetina*; N = *Nigronia*.

| | LEAVES | CBOM | FBOM | UFBOM | CPOM | FPOM | UFPOM | Fontinalis | LEM | CLAD | VAUC | PERI |
|---|---|---|---|---|---|---|---|---|---|---|---|---|
| *16:0 O* | 29.7 | 23.1 | 21.8 | 29.0 | 20.5 | 28.1 | 26.8 | 31.9 | 32.7 | 22.5 | 27.1 | 32.5 |
| *16:0 S* | 28.4 | 21.0 | 22.5 | 30.1 | 19.2 | 28.6 | 29.7 | 34.0 | 39.1 | 17.9 | | 28.8 |
| *16:1 O* | 1.2 | 0.8 | 0.5 | 4.3 | 0.0 | 2.5 | 2.6 | 3.3 | 1.6 | 3.5 | 1.4 | 0.0 |
| *16:1 S* | 0.0 | 2.0 | 0.8 | 5.7 | 0.5 | 0.9 | 0.5 | 3.0 | 1.2 | 5.5 | | 3.3 |
| ***16:1ω7 O*** | **5.4** | **7.6** | **13.6** | **18.3** | **6.0** | **10.9** | **13.5** | **15.3** | **3.9** | **10.5** | **34.6** | **20.8** |
| ***16:1ω7 S*** | **4.7** | **5.7** | **12.4** | **13.5** | **5.8** | **10.2** | **9.3** | **6.4** | **3.2** | **7.0** | | **12.4** |
| *16:2ω4 O* | 0.1 | 0.4 | 0.6 | 0.0 | 0.4 | 0.0 | 0.5 | 0.0 | 0.0 | 1.1 | 0.0 | 0.5 |
| *16:2ω4 S* | 0.0 | 0.5 | 1.2 | 0.0 | 1.3 | 0.0 | 0.2 | 0.0 | 0.0 | 0.9 | | 0.3 |
| *16:3ω4 O* | 0.0 | 0.0 | 0.0 | 0.0 | 0.0 | 0.0 | 0.0 | 0.0 | 0.0 | 1.9 | 0.0 | 0.0 |
| *16:3ω4 S* | 0.0 | 0.0 | 0.0 | 0.0 | 0.0 | 0.0 | 0.0 | 0.0 | 0.0 | 2.9 | | 0.1 |
| ***16PUFA a O*** | **0.0** | **0.2** | **0.6** | **0.8** | **0.3** | **0.0** | **0.1** | **1.6** | **0.2** | **1.6** | **4.4** | **1.9** |
| ***16PUFA a S*** | **0.0** | **0.1** | **0.5** | **0.0** | **1.0** | **0.0** | **0.0** | **0.7** | **0.0** | **1.0** | | **0.7** |
| ***16PUFAb O*** | **0.0** | **0.0** | **0.0** | **0.2** | **0.6** | **0.0** | **0.0** | **1.3** | **0.0** | **6.9** | **0.7** | **0.5** |
| ***16PUFAb S*** | **0.0** | **0.3** | **0.4** | **0.0** | **0.7** | **0.0** | **0.0** | **0.8** | **0.0** | **8.8** | | **1.2** |
| *18:0 O* | 6.1 | 4.9 | 5.9 | 5.4 | 7.0 | 10.6 | 7.0 | 1.4 | 1.5 | 0.6 | 0.8 | 3.0 |
| *18:0 S* | 4.5 | 4.2 | 5.0 | 6.8 | 5.6 | 10.4 | 6.6 | 1.8 | 1.3 | 0.8 | | 4.8 |
| ***18:1ω7 O*** | **3.5** | **7.2** | **6.2** | **6.7** | **2.6** | **4.9** | **6.2** | **1.6** | **1.6** | **2.1** | **0.4** | **5.0** |
| ***18:1ω7 S*** | **3.6** | **5.5** | **6.9** | **8.1** | **2.8** | **4.7** | **6.1** | **2.4** | **2.5** | **3.6** | | **8.0** |
| ***18:1ω9 O*** | **15.8** | **19.2** | **10.6** | **7.6** | **11.1** | **11.6** | **10.9** | **1.9** | **23.2** | **10.1** | **2.1** | **5.0** |
| ***18:1ω9 S*** | **11.1** | **10.9** | **10.8** | **9.7** | **7.7** | **11.0** | **11.1** | **2.2** | **15.5** | **10.2** | | **8.0** |
| ***ΣLCFA O*** | **5.5** | **4.7** | **7.2** | **2.8** | **4.8** | **6.3** | **6.3** | **1.5** | **0.8** | **0.0** | **0.9** | **0.4** |
| ***ΣLCFA S*** | **3.8** | **14.5** | **7.4** | **1.8** | **5.3** | **11.1** | **7.8** | **0.0** | **0.1** | **0.0** | | **1.0** |

**Table S1:** Simplified fatty acid profiles of food sources of the Cross River not represented in paper Figures. An O next to the fatty acid represents content in the open section of the river, whereas an S represents content in the shaded section. Values are mean percent fatty acid content. LEM = *Lemanea*; CLAD = *Cladophora*; VAUC = *Vaucheria*; PERI = periphyton; ΣLCFA = Sum of all long chain saturated or unsaturated fatty acids, markers of terrestrial matter (20:0;20:2;21:0;22:0;22:2;23:0;24:0). Some FAs that were not present have been excluded from table to simplify view.

| | Primary Consumers | | | | | | Predators | | |
|---|---|---|---|---|---|---|---|---|---|
| | **Hydropsyche** | **Epeorus** | **Isonychia** | **Stenonema** | **Glossosoma** | **Psephenus** | **Rhyacophila** | **Paragnetina** | **Nigronia** |
| 16:0 O | 17.4 | 21.3 | 19.7 | 18.1 | 21.3 | 20.5 | 21.2 | 16.9 | 13.1 |
| 16:0 S | 14.0 | 14.2 | 20.9 | 15.1 | 18.0 | 21.4 | 20.2 | 16.4 | 14.8 |
| **16:1ω7 O** | **10.8** | **11.4** | **9.4** | **13.2** | **10.5** | **9.1** | **14.1** | **9.3** | **8.5** |
| **16:1ω7 S** | **6.8** | **5.6** | **9.7** | **7.8** | **5.9** | **8.6** | **15.3** | **8.9** | **9.6** |
| 16:2ω4 O | 0.2 | 0.0 | 0.0 | 0.1 | 0.0 | 0.0 | 0.2 | 0.1 | 0.0 |
| 16:2ω4 S | 0.0 | 0.0 | 0.0 | 0.1 | 0.0 | 0.0 | 0.1 | 0.0 | 0.2 |
| 16:3ω4 O | 0.0 | 0.0 | 0.0 | 0.0 | 0.5 | 0.3 | 0.2 | 0.0 | 0.0 |
| 16:3ω4 S | 0.0 | 0.0 | 0.0 | 0.0 | 0.7 | 0.7 | 0.3 | 0.0 | 0.0 |
| **16PUFA a O** | **1.6** | **2.9** | **1.3** | **1.5** | **4.0** | **3.0** | **2.1** | **1.8** | **0.9** |
| **16PUFA a S** | **1.2** | **1.0** | **1.3** | **0.9** | **3.9** | **2.3** | **2.4** | **1.6** | **1.4** |
| **16PUFAb O** | **0.5** | **0.6** | **0.0** | **0.2** | **4.6** | **2.4** | **0.2** | **0.5** | **0.8** |
| **16PUFAb S** | **0.7** | **0.1** | **0.0** | **0.0** | **3.1** | **1.3** | **0.2** | **0.2** | **0.2** |
| 18:0 O | 6.1 | 5.7 | 5.7 | 3.9 | 3.9 | 2.8 | 4.9 | 6.6 | 9.4 |
| 18:0 S | 4.0 | 7.9 | 4.9 | 6.7 | 3.7 | 3.3 | 5.5 | 7.0 | 8.6 |
| **18:1ω7 O** | **2.7** | **7.0** | **8.0** | **12.9** | **2.2** | **3.3** | **3.3** | **3.6** | **5.2** |
| **18:1ω7 S** | **1.8** | **8.4** | **7.4** | **12.4** | **2.4** | **3.5** | **3.6** | **4.9** | **4.5** |
| **18:1ω9 O** | **13.5** | **7.7** | **13.2** | **7.2** | **6.0** | **16.0** | **16.3** | **18.5** | **17.0** |
| **18:1ω9 S** | **13.0** | **8.9** | **14.2** | **10.2** | **7.0** | **17.3** | **15.7** | **17.5** | **17.3** |
| ***ΣLCFA O*** | **0.5** | **0.2** | **0.1** | **0.0** | **0.2** | **0.0** | **0.6** | **0.3** | **1.4** |
| ***ΣLCFA S*** | **0.1** | **0.0** | **0.0** | **0.0** | **0.1** | **0.0** | **0.8** | **0.9** | **1.1** |

**Table S2:** Simplified fatty acid profiles of primary consumers and predators of the Cross River not represented in paper Figures. An O next to the fatty acid represents content in the open section of the river, whereas an S represents content in the shaded section. Values are mean percent fatty acid content. ΣLCFA = Sum of all long chain saturated or monounsaturated fatty acids, markers of terrestrial matter (20:0:20:2:21:0:22:0:22:2:23:0:24:0).
